# Inhibition of MLKL impairs abdominal aortic aneurysm development by attenuating smooth muscle cell necroptosis

**DOI:** 10.1101/2022.11.24.517638

**Authors:** Harshal N Nemade, Dennis Mehrkens, Hannah Sophia Lottermoser, Zeynep Ece Yilmaz, Patrick Schelemei, Felix Ruben Picard, Simon Geißen, Gülsah Fülgen Schwab, Friedrich Felix Hoyer, Henning Guthoff, Alexander Hof, Felix Sebastian Nettersheim, Agapios Sachinidis, Holger Winkels, Stefan Baldus, Manolis Pasparakis, Matti Adam, Martin Mollenhauer

## Abstract

**Background:** Receptor-interacting serine/threonine-protein kinase 1 and 3 (RIPK1 and RIPK3) dependent cell death has been identified as a crucial mediator of abdominal aortic aneurysm (AAA) development. RIPK3 mediates phosphorylation of Mixed lineage kinase domain like pseudokinase (MLKL) thereby inducing its oligomerization and translocation to the cell membrane. Given the dual role of RIPKs being involved in necroptosis as well as in apoptosis induction, the specific role of MLKL-induced necroptotic cell death in AAA remains unclear.

**Methods:** We monitored elastase-perfusion (PPE) induced progression of AAA in C57BL/6N (WT), RIPK1 kinase-inactive (*Ripk1*^*D138N/D138N*^), MLKL knockout (*Mlkl^−/−^*) and MLKL phospho-deficient (*Mlkl*^*AA*^) mice by ultrasound measurements, histological analyses and bulk mRNA sequencing to assess structural and molecular aortic changes. Bone marrow transplantations in WT and *Mlkl*^*AA*^ mice were utilized to dissect the role of MLKL in smooth muscle cells (SMCs) and myeloid cells in AAA development. MLKL expressing human SMCs were generated to investigate necroptosis-induced proinflammatory cytokine secretion and subsequent polymorphonuclear neutrophil (PMN) migration and activation in vitro.

**Results:** Ultrasound analysis showed that ~70% of the WT animals developed PPE induced-AAA with significant aortic structural alterations and enhanced myeloid cell infiltration. In contrast, *Ripk1*^*D138N/D138N*^, *Mlkl*^*AA*^, and *Mlkl^−/−^* mice were protected from AAA. This protection was associated with reduced adverse extracellular matrix (ECM) remodeling and leukocyte infiltration. MLKL deficiency was associated with a significant downregulation of genes involved in fibrinolysis, anti-inflammatory response, immune response and complement activation in aortic tissue in AAA. Bone marrow transplantation studies showed the lack of MLKL in SMCs to be the main driver of AAA protection. Proinflammatory cytokine secretion was elevated in necroptosis induced SMCs and resulted in a significant accumulation and activation of PMN.

**Conclusions:** Overall, these findings indicate that MLKL-induced necroptotic SMC death and subsequent proinflammatory leukocyte activation play a causative role in AAA development and suggest that pharmacological inhibition of MLKL may represent a promising treatment strategy for AAA disease.

## Introduction

Abdominal aortic aneurysm (AAA) formation is a common pathology in western countries with an incidence of around 2.5%.^1^ AAA disease is associated with aortic complications, including rupture or dissection, which occur in 5% of all patients and have a poor prognosis.^2^ Therefore development of therapeutic options to inhibit AAA formation and progression is of high clinical importance. Loss of smooth muscle cells (SMC) in the tunica media of the aortic wall represents a central event in the development of AAA and apoptotic cell death has been early identified as a critical mechanism of SMC degradation in this disease.^3^ Apoptosis is a well-characterized form of regulated cell death orchestrated by a family of cysteine proteases known as caspases.^4^ Apoptosis was considered for many years the only type of regulated cell death. However, the recent discovery of molecularly controlled pathways of lytic cell death such as necroptosis revealed that also necrotic cell death, which was initially suggested to represent passive, uncontrolled and ‘accidental’, can be regulated.^5^ Recent studies identified necroptosis as a regulated necrotic cell death process that is induced by receptor-interacting kinases 3 (RIPK3) and its substrate mixed lineage kinase-like protein (MLKL).^6^ Necroptosis is triggered by various stimuli such as toll-like receptor (TLR) activation, interferon-gamma (IFN-γ), or intracellular DNA and RNA recognition, with tumor necrosis factor alpha (TNF-α) signaling being the most studied mechanism until now. Activation of the TNF-α receptor results in trans- and autophosphorylation of RIPK1 which interacts with RIPK3 to induce the assembly of the necrosome complex that facilitates the subsequent phosphorylation of MLKL by RIPK3 at serine S345 and S347.^7,8^ MLKL phosphorylation exposes its N-terminal domain inducing trimerization and translocation of MLKL to plasma membrane thereby “executing” necroptotic cell death by inducing Ca^2+^ influx and cellular lysis.^9^ In contrast to apoptosis that is considered as non-inflammatory type of cell death, necroptosis serves as a potent inducer of inflammation and has been linked to inflammatory diseases of the skin and the gut, as well as to chronic liver inflammation, neurodegenerative pathologies and cancer.^10^

The formation and progression of AAA disease is a highly inflammatory process. Progressive dilation of the aorta is associated with the recruitment and activation of leukocytes, such as polymorphonuclear neutrophils (PMN) and macrophages. Subsequent inflammatory activation of myeloid cells leads to production of reactive oxygen species (ROS) and pro-inflammatory cytokines causing subsequent inflammatory cell infiltration. These processes finally elicit degradation of the extracellular matrix (ECM) by matrix metalloprotease (MMP) activation which further drives AAA progression. Necroptosis induced membrane permeabilization rapidly releases cellular components and damage-associated molecular patterns (DAMPs),^11^ which are recognized by resident aortic macrophages. In turn, this leads to the formation of the NLRP3 inflammasome triggering further production of proinflammatory cytokines and chemokines including TNF-α and IL-1β, a hallmark of AAA development.^12,13^

Although depletion of PMN^14^ or IL-1β^15^ inhibition showed promising results in animal models and clinical trials have confirmed the efficacy of anti-inflammatory therapies in atherosclerotic cardiovascular diseases (CANTOS^16^, COLCOT^17^), no targeted therapies for AAA disease are available so far.^18^ Pharmacological inhibition of RIPK1 and genetic loss of RIPK3 ameliorates AAA progression in mice pointing to cell death as a potential target in AAA disease treatment.^19,20^ Nonetheless, given the dual role of both kinases being involved in necroptosis and in apoptosis, the specific role of necroptotic cell death in AAA remains unclear. Whether necroptotic stimuli in the early phase of AAA formation activate leukocytes, and/or the recruitment of leukocytes drives the extent of aneurysm formation by induction of additional necroptosis remains to be elucidated.

## Results

### Loss of MLKL attenuates aneurysm formation in mice

To investigate the role of MLKL induced-necroptosis in the development and progression of AAA disease 10-to 14-week old male WT, Mlkl knockout (*Mlkl^−/−^*), RIPK1 kinase-inactive (*Ripk1*^*D138N/D138N*^) and MLKL phospho mutated (*Mlkl*^*AA*^) mice were subjected to PPE surgery. *Mlkl^−/−^*, *Mlkl*^*AA*^ and, in part, *Ripk1*^*D138N/D138N*^ animals showed an attenuation of aortic diameter increase as assessed by ultrasound analysis compared to WT mice (Fig. 1A and 1B). Interestingly, *Ripk1*^*D138N/D138N*^, *Mlkl*^*AA*^, and *Mlkl^−/−^* mice were protected from AAA induction, whereas ~70% of WT animals developed PPE-induced AAA (Supplemental Fig. S1A). Masson’s trichrome staining (MTS) and elastin fluorescence imaging showed that aortic structural alterations, collagen degradation and elastin disarray were ameliorated in *Mlkl^−/−^* and *Mlkl*^*AA*^ animals, however less significant improvements in aortic wall structural components were observed in *Ripk1*^*D138N/D138N*^ mice 3 and 28 days after AAA induction (Fig. 1C and 1D, MTS and elastin grade definition is shown in Supplemental Fig. S7). TEM micrographs demonstrated decreased EC and SMC cell volume, loss of SMC membrane integrity and disarranged collagen fibers in WT animals subjected to AAA. Theses alterations were strongly attenuated in *Mlkl^−/−^*, *Mlkl*^*AA*^ and, in part, in *Ripk1*^*D138N/D138N*^ animals (Fig. 1E upper panel). Furthermore, WT animals developed disarranged and fragmented collagen fiber structure post-PPE, which was not detectable in *Mlkl^−/−^* and *Mlkl*^*AA*^ mice (Fig. 1E lower panel, Supplementary Fig. S1B). Interestingly, *Ripk1*^*D138N/D138N*^ animals showed marked changes in SMC morphology, elastin degradation and loss of ECs as compared to WT.

**Figure 1.**
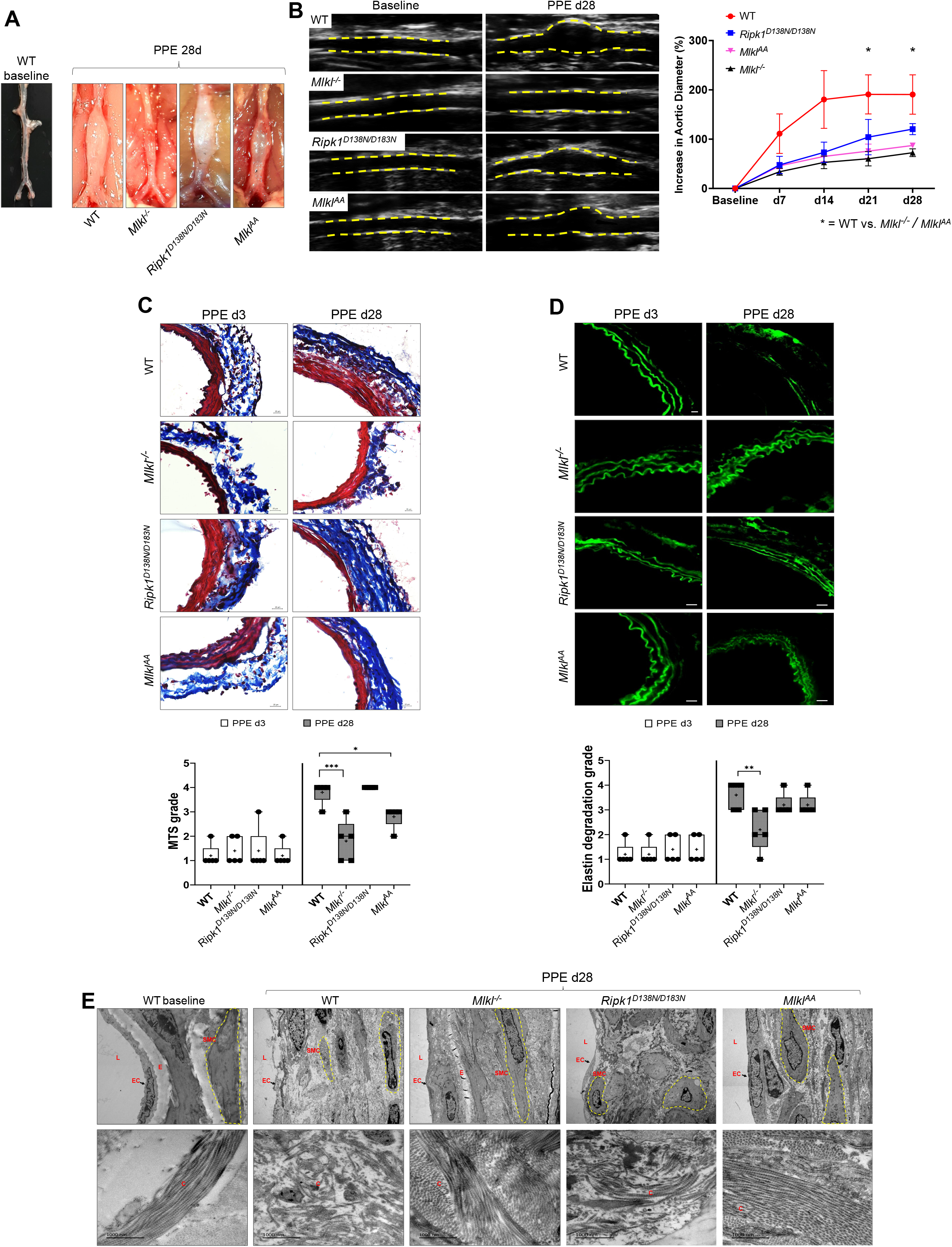
Necroptotic cell death is crucial for aortic dilation and vascular remodeling in AAA. (**A**) Representative macrographs of abdominal aorta at baseline and 28 days post PPE (scale bar = 1 mm). (**B**) Representative B-mode ultrasound images of abdominal aorta at baseline and 28 days post PPE. Yellow lines indicate vascular wall. Assessment of aortic diameter represented as percent increase to baseline diameter (n = 5 per group). (**C**) Representative images of abdominal aortic sections stained with Masson’s Trichrome Staining (MTS) (scale bars = 20 μm) and quantification of aortic collagen content (red) by MTS grade analyses (n = 5 per group). (**D**) Representative images of elastin autofluorescence of abdominal aortic sections (green, scale bars = 20 μm) and quantification of elastin strand breaks by elastin degradation grade analyses (n = 5 per group). (**E**) Representative transmission electron microscopy (TEM) images of abdominal aortic sections at baseline and 28 days post AAA displaying smooth muscle cells (SMC) morphology, endothelial cell (EC) abundance and elastin (E) degradation (upper panel) and collagen (C) structure and fiber arrangement (lower panel; L = Lumen; scale bars = 1000 nm). Data are expressed as mean ± SEM. Statistical significance was determined by one-way ANOVA with Tukey’s multiple comparisons test for. Wildtype (WT), MLKL deficient (*Mlkl^−/−^*), RIP1 kinase dead (*Ripk1*^*D138N/D183N*^), MLKL mutated (*Mlkl*^*AA*^). * = *P<0*.*05*, ** = *P<0*.*01*, *** = *P<0*.*005*, **** = *P<0*.*001*).

### Loss of MLKL abrogates the infiltration of immune cells in PPE-induced AAA lesions

To determine whether the inhibition of necroptosis abrogates the infiltration of immune cells, the abdominal aorta of mice was harvested and analyzed by immunofluorescence staining, histology and flow cytometry. At baseline conditions no Ly6G^+^ PMN or CD68^+^ macrophages could be detected within the aortae of WT animals (Supplemental Fig. S2A). 3 days post-PPE, WT animals showed marked infiltration of Ly6G^+^ PMN in AAA lesions, whereas *Mlkl^−/−^*, *Ripk1*^*D138N/D138N*^ and *Mlkl*^*AA*^ mice harbored significantly less Ly6G^+^ cells in the tunica media (Fig. 2A and 2B). Immunofluorescence stainings revealed enhanced CD68^+^ macrophages numbers within the tunica media (Fig. 2C and 2D) which was attenuated in *Mlkl^−/−^*, *Ripk1*^*D138N/D138N*^ and *Mlkl*^*AA*^ mice. Interestingly, the location of PMN and macrophages coincided with the loss of alpha smooth muscle cell actin (aSMA) signal within the tunica media indicating enhanced SMC loss at the site of inflammatory cell infiltration. To confirm whether reduced leukocyte infiltration by MLKL deficiency is a long-term effect, we further investigated aortae of all genotypes 28 days after PPE. No Ly6G^+^ signal could be detected whereas CD68^+^ macrophage infiltration was significantly attenuated in the tunica media of *Mlkl^−/−^*, *Ripk1*^*D138N/D138N*^ and *Mlkl*^*AA*^ animals (Supplementary Fig. S2B and S2C).

**Figure 2.**
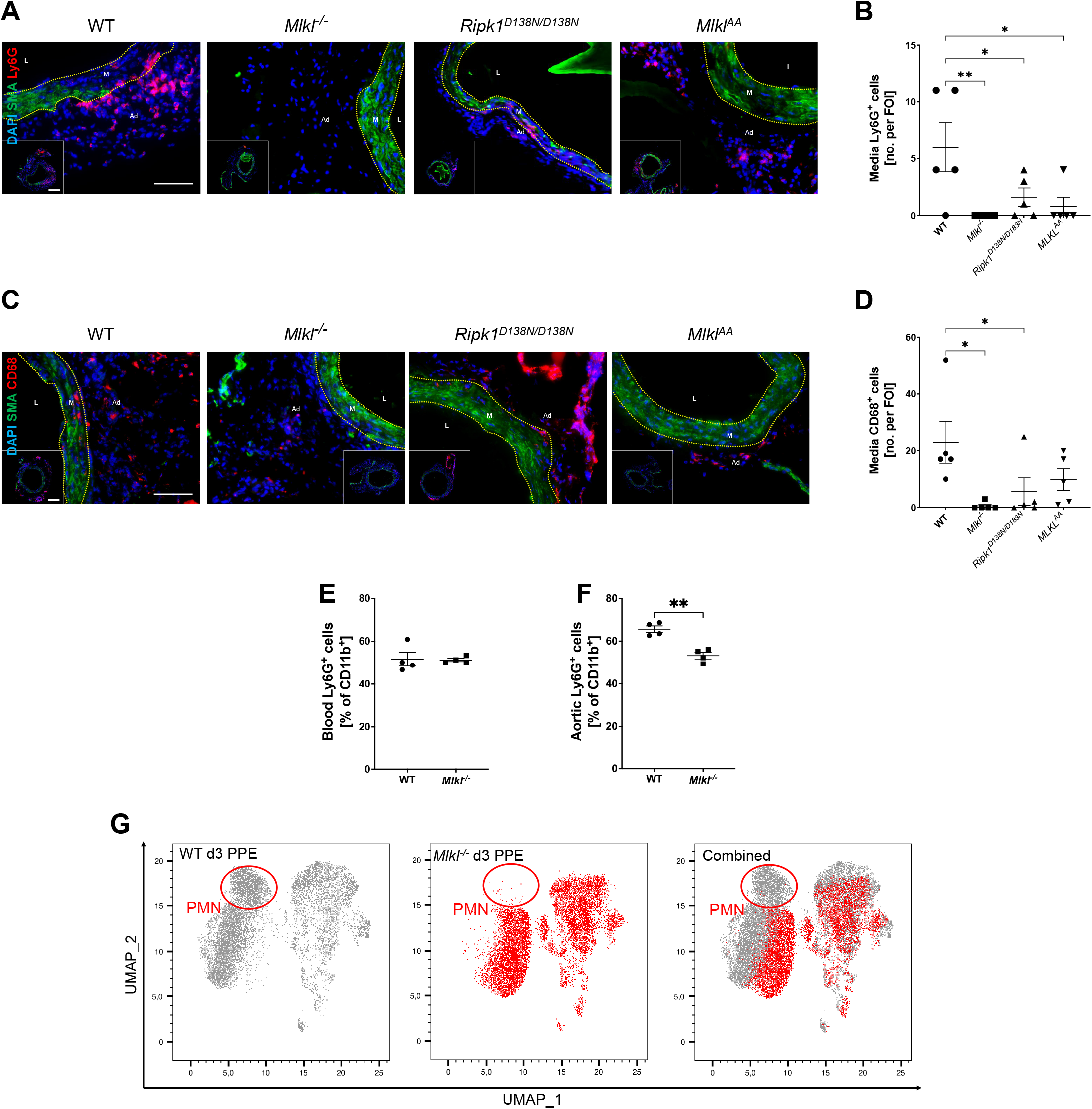
Lack of MLKL reduces proinflammatory immune cell infiltration during early stages of AAA. (**A**) Representative confocal aortic fluorescence images of the smooth muscle cell marker a-SMA (green) and of the PMN marker Ly6G (red) 3 days post PPE and quantification of aortic Ly6G^+^ cells within (**B**) the tunica media (n = 5 per group). (**C**) Representative confocal aortic fluorescence images of the smooth muscle cell marker a-SMA (green) and the macrophage marker CD68 (red, large scale bar = 100μm, small scale bar = 20μm; vascular lumen (L), tunica media (M), tunica adventitia (Ad)) and quantification of aortic CD68^+^ cells within (**D**) the tunica media (n = 5 per group). (**E**) Flow cytometric analyses of Ly6G^+^ cells in the blood and (**F**) in the abdominal aorta 3 days after PPE (n = 4 per group). (**G**) Uniform manifold approximation and projection (UMAP) dimensional reduction plot of WT and *Mlkl^−/−^* aortae 3 days post-PPE. Data are expressed as mean ± SEM. Statistical significance was determined by one-way ANOVA with Tukey’s multiple comparisons test. Wildtype (WT), MLKL deficient (*Mlkl^−/−^*), RIP1 kinase dead (*Ripk1*^*D138N/D183N*^), MLKL mutated (*Mlkl*^*AA*^). * = *P<0*.*05*, ** = *P<0*.*01*, *** = *P<0*.*005*).

To further investigate the effect of MLKL in aortic inflammation during AAA development, we performed flow cytometry analyses of the blood and aorta of WT and *Mlkl^−/−^* mice 3 days post PPE. The detailed gating strategy is shown in Supplementary Fig. S2D. We could identify 16 unique cell clusters based on the cell specific marker expression levels (Supplementary Fig. S2E). PMN blood count was equal in WT and *Mlkl^−/−^* animals (Fig. 2E) whereas total aortic PMN infiltration was attenuated in *Mlkl^−/−^* animals compared to WT (Fig. 2F). Further UMAP analysis showed WT specific PMN populations which were either significantly reduced or completely absent in the aortae of *Mlkl^−/−^* mice (Fig. 2G, red circle).

### Necroptosis deficiency downregulates inflammation and fibrinolysis related genes in AAA

Abdominal aortic tissue samples were collected from WT (WT 3d PPE) animals 3 days post PPE and bulk mRNAseq was performed. Unoperated WT animals were used as baseline control (WT bl). The mRNAseq analysis identified 6954 differently expressed genes (DEGs) (≥1 log2FC, *p*>0.05) in WT animals 3 days post-PPE compared to WT bl animals consisting of 3889 upregulated and 3065 downregulated genes (Fig. 3A and 3B). A majority of the upregulated genes were enriched in cytokine production, immune and inflammation response pathways, leukocyte-, neutrophil- and macrophage-migration related GO terms, comprising for example *Cxcl3* (log2FC =15.65), *Cxcl5* (log2FC =14.26), *Il1a* (log2FC =12.12), *Il1b* (log2FC =11.78), *Ccl3* (log2FC =10.69), *Il6* (log2FC =10.55), and *Csf3* (log2FC =10.25) (Fig. 3C and 3D, Supplemental Fig. S3A) whereas most downregulated genes were enriched in muscle contraction, smooth muscle cell contraction and cation transport related GO terms for example *Myh6* (log2FC = −6.85), *Slc2a5* (log2FC = −6.04), *Mstn* (log2FC = −5.88), *Mb* (log2FC = −5.25), *Myl1* (log2FC = −4.98), *Myh6* (log2FC = −6.85), and *Cacna1s* (log2FC = −3.04) (Fig. 3E and 3F, Supplemental Fig. S3B). These findings correlate with our observed increase in aortic Ly6G^+^ and CD68^+^ cells. In addition to inflammation related processes, genes involved in fibrinolysis including *Hrg, Fga, Fgg, F2, Fgb, Apoh* and *Plg* were significantly upregulated in WT d3 PPE animals (Supplemental Fig. S3C).

**Figure 3.**
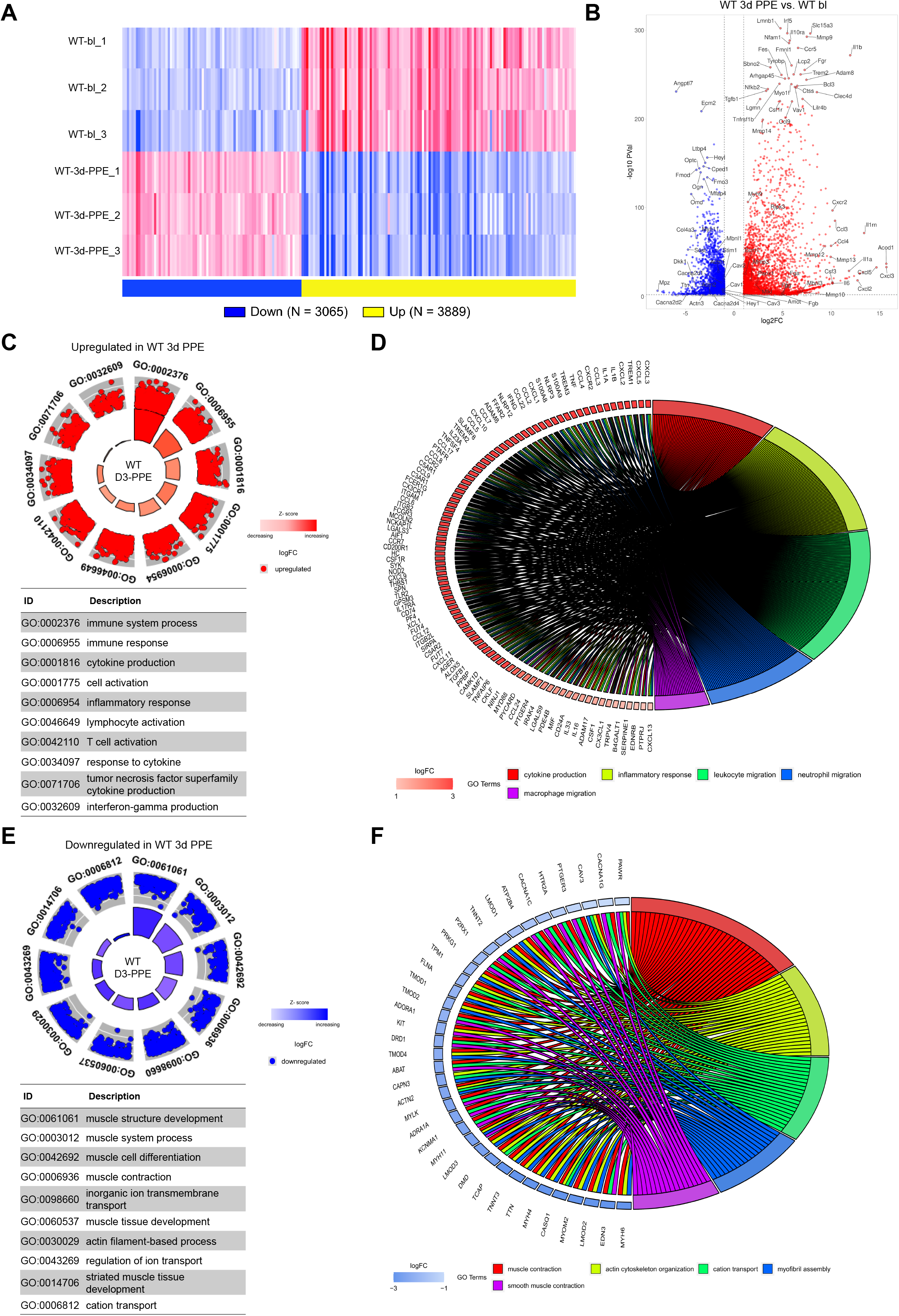
AAA aortae acquire an inflammatory transcriptional phenotype. Aorta from WT baseline (bl) and WT 3 days post PPE were harvested for bulk transcriptome analysis (n = 3 per group). (**A**) Heatmap visualizing the significantly differentially expressed (DE) genes between WT bl and WT d3 PPE. (**B**) Volcano plot showing the sufficiently up- (red) and downregulated (blue) DE genes between WT bl and WT 3d PPE mice. The top DE genes are labelled. (**C)** The unique differentially expressed genes were subjected to pathway analysis. GOcircle plot display scatter plots of upregulated genes with their log fold change (logFC) and the most statistically significant GO terms (table). Inner bars indicate the significance of the corresponding GO terms (-log10-adjusted P-value), and the color corresponds to the enrichment Z-score calculated by GOplot function in R. (**D**) GOChord plot showing assignment of upregulated genes to their respective enriched GO categories (BP) according to their fold change. (**E**) GOcircle plot display scatter plots of downregulated genes with their log fold change (logFC) and the most statistically significant GO terms. Inner bars indicate the significance of the corresponding GO terms (-log10-adjusted P-value), and the color corresponds to the enrichment Z-score calculated by GOplot function in R. (**F**) GOChord plot showing assignment of downregulated genes to their respective enriched GO categories (BP) according to their fold change.

Interestingly, our mRNAseq data shows 202 genes were differentially expressed in *Mlkl^−/−^* d3 PPE aortas compared to WT d3 PPE aortas (Supplemental Fig. S3E). Among them, 163 genes associated with GO terms like fibrinolysis, inflammatory response, immune response, lymphocyte and leukocyte mediated immunity and complement activation were downregulated whereas, 39 genes associated with cation transport, endocrine processes, response to stress, response to wounding and response to oxidative stress were upregulated (Fig. 4A, Supplemental Fig. S4A). In addition, we also identified 11 uniquely downregulated genes in *Mlkl^−/−^* d3 PPE animals as compared to *Mlkl*^*AA*^ d3 PPE and *Ripk1*^*D138N/D138N*^ d3 PPE animals which included known AAA and inflammation associated genes like *Ctla4*^21^, *Marco*^22,23^, *Csf3*^24^, *Chil1*^25^ and *Olfm4*^26,27^ (Supplemental Fig. S3D). Downregulation in these genes further support our initial observations of significantly reduced inflammatory cell infiltration in aortic lesions of *Mlkl^−/−^* animals post PPE.

**Figure 4.**
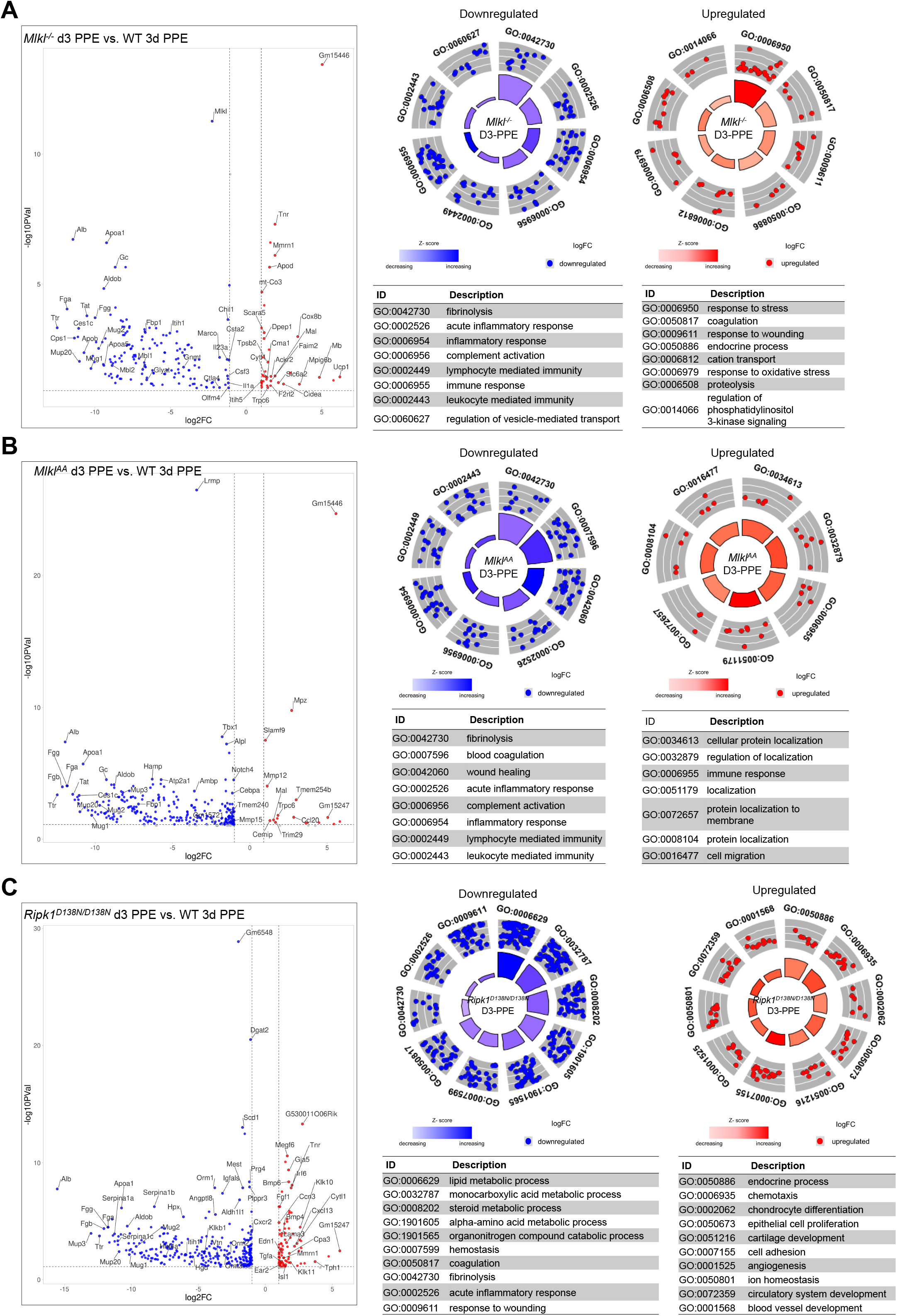
Necroptosis deficiency downregulates inflammation and fibrinolysis pathway related genes in AAA. Volcano plot showing the sufficiently up- (red) and downregulated (blue) DE genes between (**A**) *Mlkl^−/−^* d3 PPE vs. WT 3d PPE, (**B**) *Mlkl*^*AA*^ d3 PPE vs. WT 3d PPE and (**C**) *Ripk1*^*D138N/D138N*^ d3 PPE vs. WT 3d PPE mice with unique differentially expressed genes subjected to pathway analysis, respectively. GOcircle plot display scatter plots of upregulated genes with their log fold change (logFC) and the most statistically significant GO terms (table). Inner bars indicate the significance of the corresponding GO terms (-log10-adjusted P-value), and the color corresponds to the enrichment Z-score calculated by GOplot function in R.

We also performed mRNAseq analysis with aortic tissue harvested from *Mlkl*^*AA*^ (*Mlkl*^*AA*^ d3 PPE) and *Ripk1*^*D138N/D138N*^ (*Ripk1*^*D138N/D138N*^ d3 PPE). When compared to WT d3 PPE, we found 294 and 411 DEGs in *Mlkl*^*AA*^ d3 PPE and *Ripk1*^*D138N/D138N*^ d3 PPE animals respectively (Supplemental Fig. S3F and S3G). DEGs were associated with downregulation of inflammatory responses although we found some genes upregulated in immune response and chemotaxis related GO terms (Fig. 4B and 4C and Supplemental Fig. 4B and 4C) and fibrinolysis pathway related genes were significantly downregulated in both *Mlkl*^*AA*^ and *Ripk1*^*D138N/D138N*^ animals (Supplemental Fig. S3C). Taken together, inhibition of necroptosis not only leads to downregulation in AAA mediated inflammatory signaling but also in fibrinolysis. Although the link between necroptosis and fibrinolysis has not been described so far, this data point towards the potential molecular mechanisms associated with necroptosis and its role in AAA development.

### Lack of MLKL in abdominal aortic SMCs is protective against AAA development

Next, we investigated whether loss of MLKL in SMC or hematopoietic cells contributes to AAA protection and attenuated aortic leukocyte infiltration. In a bone marrow (BM) transplantation study (Supplemental Fig. S5A) irradiated recipient WT mice were reconstituted with *Mlkl*^*AA*^-BM (WT^MLKLAA-BM^) and irradiated *Mlkl*^*AA*^ recipient mice were reconstituted with WT-BM (Mlkl^AA+WT-BM^) generating SMC (*WT*)/ macrophage (*Mlkl*^*AA*^) and SMC (*Mlkl*^*AA*^)/ macrophage (WT) chimera mice. WT mice transplanted with WT-BM (WT^WT-BM^) were used as controls. WT^WT-BM^ animals developed aneurysms with significant increase in aortic diameter as indicated by ultrasound investigations. WT^MLKLAA-BM^ animals showed increased abdominal aortic diameter in comparison to Mlkl^AA+WT-BM^ animals which were protected from AAA development (Fig. 5A and 5B). Histological stainings confirmed that Mlkl^AA+WT-BM^ animals have stable aortic wall structures post PPE, whereas both other groups showed significantly elevated remodeling processes with disarranged collagen (Fig. 5C and Supplemental Fig. S5B upper panel), loss of media boundary, and increased elastin strand breaks (Fig. 5E and Supplemental Fig. S5B lower panel). We hypothesized that lack of MLKL would prevent SMC necroptosis and in turn reduce proinflammatory cell infiltration in Mlkl^AA+WT-BM^ aortae. Indeed, we observed significantly reduced Ly6G^+^ cells in the abdominal aorta of Mlkl^AA+WT-BM^ animals (Fig. 5E and Supplemental Fig. S5C upper panel) with no significant reduction in CD68^+^ cell number (Fig. 5F and Supplemental Fig. S5C lower panel).

**Figure 5.**
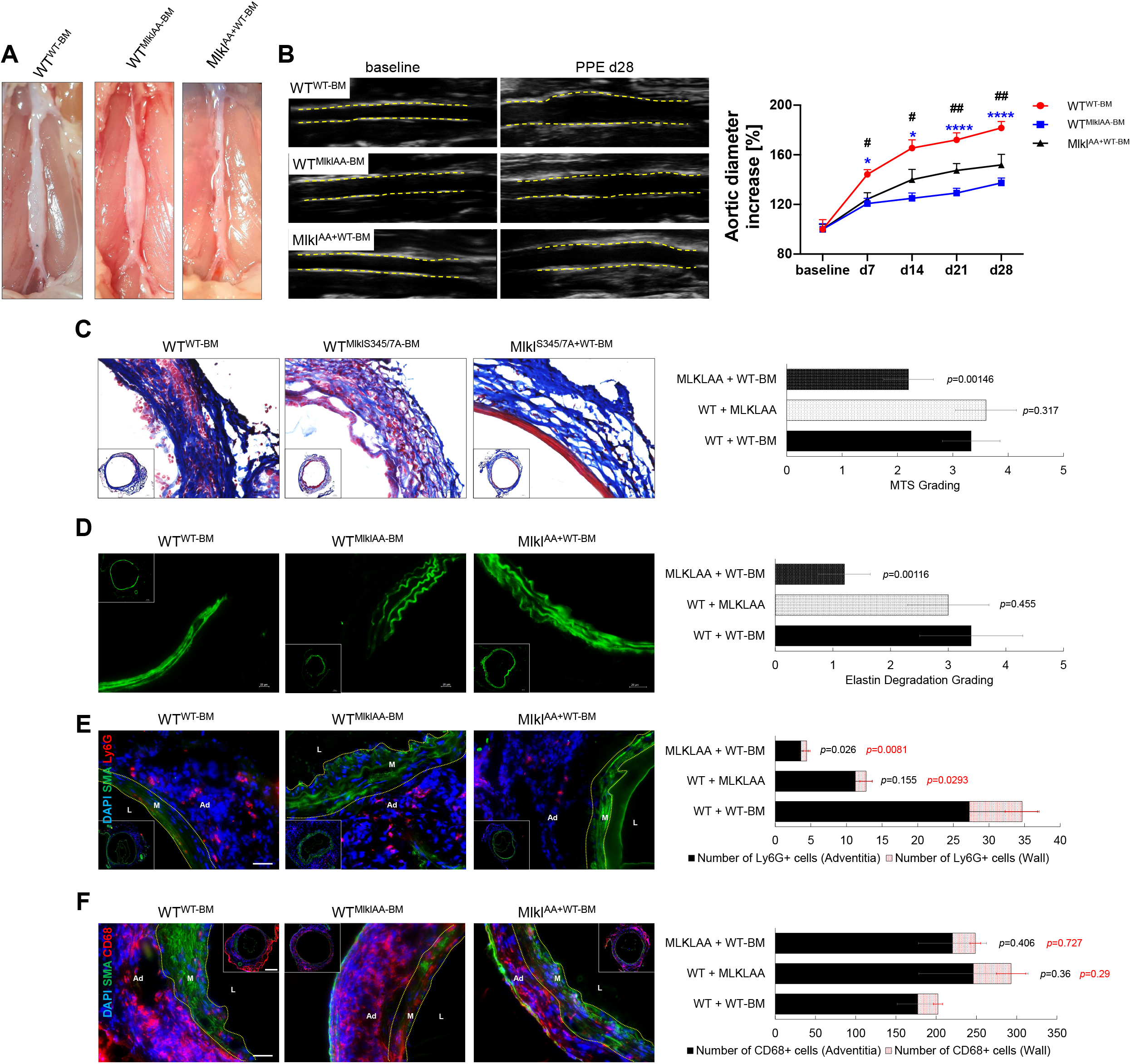
MLKL deficiency in aortic SMC is protective against AAA development. (**A**) Representative macrographs of abdominal aorta 28 days post PPE. (**B**) Representative B-mode ultrasound images of abdominal aorta at baseline and 28 days post PPE and quantification of the aortic diameter. Yellow lines indicate the vascular wall (n = 5 per group). (**C**) Representative images of Masson’s Trichrome stained (MTS) abdominal aortic sections (scale bars = 100μm and 20μm) and quantification of aortic collagen content (red) by MTS grade analyses (n = 5 / 5 / 6). (**D**) Representative images of elastin autofluorescence of abdominal aortic sections (green) and quantification of elastin strand breaks by elastin degradation grade analyses (n = 5 for all groups). (**E**) Representative confocal aortic fluorescence images of the smooth muscle cell marker a-SMA (green) and the PMN marker Ly6G (red) and quantification of total (black) Ly6G^+^ cells and Ly6G^+^ cells within the tunica media (red) (n = 5 for all groups) and (**F**) the macrophage marker CD68 (red) and quantification of total (black) CD68^+^ cells and CD68^+^ cells within the tunica media (red) (n = 5 per all groups). Scale bars = 100μm and 20μm. Vascular lumen (L), tunica media (M), tunica adventitia (Ad). Data are expressed as mean ± SEM. Statistical significance was determined by one-way ANOVA with Tukey’s multiple comparisons test.

### MLKL-induced SMC death leads to release of DAMPs and promotes PMN activation

To show that MLKL-induced necroptotic SMC death is a crucial first step in the activation of PMN and subsequent AAA development, we generated two human aortic smooth muscle cells (hSMC) lines by transfecting Tet-On 3G inducible expression constructs encoding for full length human MLKL (F-hMLKL) and ‘N’ domain of human MLKL (ND-hMLKL) (aa 1-182) (Fig. 6A).^28^ Doxycycline (Dox) induced robust production of the respective MLKL mRNA detected by qRT-PCR and proteins indirectly detected via the mCherry fluorescence signal (Fig. 6B and C). ND-hMLKL isoform was 4-fold higher expressed in comparison to the F-hMLKL isoform. This difference in expression was accompanied by significantly higher SMC death in ND-hMLKL-SMC (~80%) compared to F-hMLKL-SMC (~50%). Treatment of necrosulfonamide (NSA), a well characterized MLKL inhibitor, prevented SMC death in both cell lines (Fig. 6D). Live cell imaging of transfected and induced SMCs showed membrane rupture and cell lysis indicative of typical necroptotic death (Supplemental Video V1; yellow circles). Secretome antibody array analyses of MLKL expressing SMCs confirmed a significant increase in proinflammatory cytokine secretion, e.g. *Il1a, Il1b, Mcp2, Mip3a, Nap2, Nt3* and Osteoprotegerin, whereas necroptosis inhibition by NSA attenuated cytokine release (Fig. 6E and Supplemental Fig. S6A). Given that these cytokines play a substantial role in leukocyte recruitment in AAA disease^29^, we generated a PMN-like cell line in vitro by differentiating HL60 cells (Fig. 6F) and performed co-culture studies. Indeed, MLKL expression in SMCs rapidly led to PMN attraction and phagocytosis of SMC (Fig. 6G and Supplemental Video V2), indicating MLKL induced necroptosis to be an inducer of leukocyte recruitment and activation in AAA.

**Figure 6.**
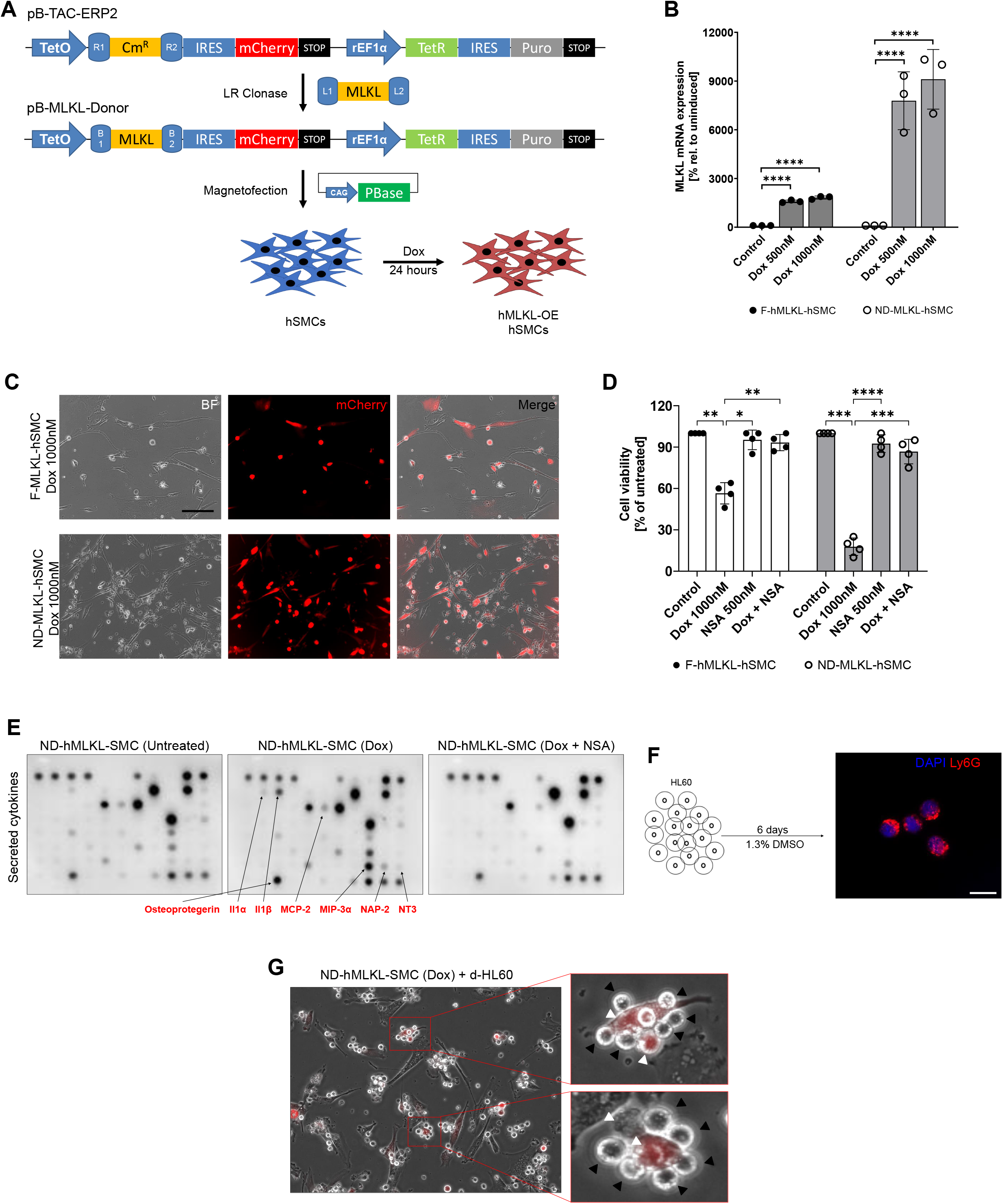
SMC necroptosis induces proinflammatory cytokine secretion and subsequent PMN activation. (**A**) Schematic of the cloning strategy to produce MLKL expressing human SMC (hSMC). The human full MLKL gene (F-hMLKL) or the ‘N’ domain of human MLKL (ND-hMLKL) was cloned into a Doxocyclin (Dox) inducible Tet-On 3G expression vector pB-TAC-ERP2. (**B**) qRT-PCR expression analyses for MLKL mRNA expression in F-hMLKL-hSMC and ND-hMLKL-hSMC (n=3 per group). (**C**) Representative fluorescence images of the mCherry signal of Dox induced F-hMLKL-hSMCs and ND-hMLKL-hSMCs. (**D**) Cell viability assay of control-, Dox, necrosulfonamide (NSA) or Dox + NSA treated F-hMLKL-hSMC and ND-hMLKL-hSMC (24h, n = 4 per group). (**E**) Representative dot blots of secretome antibody array analyses of the supernatant of control, Dox (1000nM) or Dox (1000nM) + NSA (500nM) treated ND-hMLKL-hSMCs with arrows indicating significantly changed cytokines (treatment: 24h). (**F)** Schematic and representative immunofluorescence staining for the PMN marker Ly6G of HL60 cell differentiated to mature PMN by treatment with 1.3% DMSO for 6 days. (**G**) Representative images of co-culture experiments (n = 3) with Dox induced ND-hMLKL-hSMCs (white arrow heads) and differentiated HL60 (black arrow heads) indicating accumulation of PMN around necroptotic SMCs and phagocytosis. * = *P<0*.*05*, ** = *P<0*.*01*, *** = *P<0*.*005*, **** = *P<0*.*001*.

## Discussion

We herein reveal that necroptotic cell death in SMCs is essentially involved in the pathogenesis of abdominal aortic dilation and potent mediator of vascular inflammation in AAA disease. We here provide the first evidence identifying the necroptotic executer protein MLKL as a major mediator of adverse vascular remodeling, leukocyte activation and infiltration and transcriptional alterations in AAA development. AAA is a multifactorial disease, in which loss of vascular SMCs fundamentally contributes to aortic dysfunction and ECM degeneration, ultimately cumulating in fatal aortic complications such as rupture or dissection.^30^ Furthermore, SMC-mediated paracrine effects on the tunica adventitia contribute to artery wall homeostasis with potent anti-inflammatory and anti-proteolytic properties.^31^ Hence, prevention of SMC death is of primary importance in prevention of AAA growth and aortic rupture.^32^

Given that cell death-inducing kinases, such as RIPK1 are not exclusively involved in the induction of necroptosis^33^, we herein strife to reveal the role of necroptosis on AAA disease at different points of the necroptotic signalling cascade. To achieve this goal, we utilized three different mouse models and subjected them to experimental AAA induction. MLKL deficient animals (*Mlkl^−/−^*) lack the specific necroptosis execution protein and are considered fully “necroptosis” deficient. Accordingly, phosphorylation site mutated MLKL mice (*Mlkl*^*AA*^)^34^ are not able to facilitate MLKL trimerization, translocation to and disruption of cellular membranes^34^. RIPK1 kinase-inactive animals (*Ripk1*^*D138N/D138N*^) have been shown to be protected from TNF induced necroptosis.^35^ Nonetheless, RIPK1 phosphorylation has also been linked to apoptosis induction.^36^

AAA risk, aortic dilation, ECM degradation and SMC cell death were significantly decreased in all three mouse models highlighting the prominent role of necroptotic processes in AAA. Interestingly, while *Mlkl^−/−^* and *Mlkl*^*AA*^ mice showed high protection against AAA development, *Ripk1*^*D138N/D138N*^ mice were only partially protected from aortic dilation compared to WT mice, indicating that AAA formation is in part independent of RIPK1 induced apoptosis. TEM micrographs demonstrated decreased EC and SMC cell volume, loss of SMC membrane integrity and disarranged collagen fibers in the WT animals subjected to AAA. Theses alterations were strongly attenuated in *Mlkl^−/−^*, *Mlkl*^*AA*^ and, only in part, in *Ripk1*^*D138N/D138N*^ animals which could explain partial AAA protection in this genotype.

Given the importance of inflammatory processes in AAA development, we investigated the role of necroptosis in aortic leukocyte infiltration. Necroptosis inhibition significantly reduced the number of Ly6G^+^ PMN and CD68^+^ macrophages within the tunica media and was associated with an enhanced number of SMCs compared to WT aneurysms.

Finally, our bone marrow transplantation studies clearly showed that loss of MLKL function in SMCs, reduces aortic diameter increase, stabilizes vascular integrity and significantly attenuates immune cell infiltration as compared to WT controls subjected to AAA. Mechanistically, this could be attributed to an enhanced increase of pro-inflammatory cytokines secreted by necroptotic SMCs in vitro. Necroptosis leads to rapid plasma membrane permeabilization and release of damage-associated molecular patterns (DAMPs).^11^ These DAMPs initiate immune responses through the activation of classical pattern recognition receptors (PRRs) including Toll-like receptors (TLRs), NOD-like receptors (NLRs), retinoic acid-inducible gene I (RIG-I)-like receptors (RLRs), and C-type lectin receptors (CLRs) expressed on PMN and macrophages.^37^ Furthermore, DAMPs like HMGB1, IL-1α and IL-1β can lead to further infiltration of proinflammatory immune cells.^38–40^ We here demonstrate, that MLKL induced necroptosis in SMCs leads to the secretion of pro-inflammatory cytokines, especially *Il1a, Il1b* and *Nap2* which in turn attracts and activates PMN.^41^ Thus necroptosis-related release of proinflammatory cytokines induced recruitment of inflammatory leukocytes in aneurysmatic aortic tissue.

Although bone marrow transplantation experiments indicated that necroptosis of SMCs is a major driver of AAA formation, additional effects of necroptotic cell death on endocrine myeloid cell activation could not be fully excluded. WT^MLKLAA-BM^ mice showed, at least in part, protection from aortic dilation. Other groups reported MLKL being relevant for pro-inflammatory neutrophil extracellular traps (NETs) formation, a mechanism that is also linked to AAA formation.^42,43^ Further dissection of SMC- and PMN-mediated effects in vascular diseases might therefore represent an interesting goal of future studies. Nonetheless, we showed that MLKL inhibition indeed prevents the release of proinflammatory cytokines in SMCs and reduces subsequent PMN activation which can be exploited as a therapeutic option to treat AAA.

To obtain deeper insights into the mechanistic role of necroptosis in aneurysm formation, we performed bulk mRNAseq analysis of abdominal aortae 3 days post PPE. AAA in WT animals induced upregulation of genes correlating with immune cell infiltration and vascular inflammation such as, *Cxcl3, Cxcl5, Il1a, Il1b, Ccl3, Il6, Csf3* etc., underlining the strong impact of inflammatory processes during early AAA development. Gene ontology (GO) analysis of revealed a particular enrichment of biological processes related to cytokine production, inflammatory responses, leukocyte and T cell activation. Downregulated genes correlated strongly with loss of SMCs and were enriched in GO terms like muscle contraction, actin filament-based process, cation transport, cytoskeleton organization, smooth muscle contraction and aorta development. Necroptosis deficiency in *Mlkl^−/−^*, *Mlkl*^*AA*^ and *Ripk1*^*D138N/D138N*^ lead to a significant downregulation in genes associated with leukocyte-mediated immunity and acute inflammatory immune responses, whereas genes associated with oxidative stress responses and ion transport were upregulated. Given that coagulation and fibrinolysis are known to play a vital role in the development and rupture of aneurysms ^44–47^, the downregulation of genes associated with these pathways further underlines the important role of necroptosis not only in vascular inflammation and stress responses but also in structural remodeling in the early stages of AAA.

Although our understanding of the underlying pathomechanism of AAA development and progression has greatly improved over the last decades, there is still no specific medical therapy available to slow the expansion rate of AAA^48^ and surgical interventions such as open aneurysm repair (OAR)^49^ or endovascular aneurysm repair (EVAR)^50^ are still the gold standard by improving patient survival and health-related quality-of-life^51^. In this regard, cell death pathways, in particular apoptosis and necroptosis, have been studied extensively.^52,53^ This led to the discovery of potent antiapoptotic agents, such as Q-VD-OPh, 4-PBA, and L-NIL, which are currently under clinical investigation.^54–57^ Necroptosis inhibitors targeting RIPK1 such as, necrostatin-1 and necrostatin-1s have already been shown to be effective against AAA development in mice models.^20,58,59^ Nonetheless, RIPK1s dual action in necroptosis and apoptosis regulation could explain observed off-target effects of these compounds thereby limiting their translation into an clinical approach^60^. Of note, several novel RIPK1 inhibitors like GSK’772 or GSK’547, are currently under investigation^61^. Based on our in vitro study, specific inhibition of MLKL showed strong beneficial effects in terms of reducing SMC death and reduction in secretory proinflammatory cytokine expression. Although necroptosis independent functions cannot be fully excluded^62^, targeting MLKL might be advantageous since it is considered as solely executor protein of necroptosis. In light of novel specific MLKL inhibitors currently under investigation, pharmacological MLKL inhibition might emerge as a promising treatment strategy for AAA prevention.

Taken together, this study not only dissects the critical role of MLKL-induced necroptosis in development of AAA but also identifies the therapeutic potential of targeting MLKL for AAA prevention.

## Materials and Methods

### Animal Studies

Generation of *Ripk1*^*D138N/D138N*^ mouse line is described elsewhere^35^. The generation of *Mlkl^−/−^* and *Mlkl*^*AA*^ strains will be described elsewhere (Koerner et al., in preparation). All animal studies were approved by the local animal care authorities; Landesamt für Natur, Umwelt und Verbraucherschutz Nordrhein-Westfalen (LANUV), NRW, Germany (AZ: 2018.A030 and 2016.A212) and conformed to the guidelines from Directive 2010/63/EU of the European Parliament on the protection of animals used for scientific purposes. All experiments were performed on male mice, as AAA predominantly affect men and littermates were used as controls. All mice had free access to standard laboratory diet (Altromin 1324 P Best, Altromin GmbH & Co. KG, Lage, Germany) and water. All surgical interventions were performed under isoflurane anesthesia and buprenorphine analgesia.

### PPE infusion model

To induce murine AAA, the PPE infusion was performed in 10 to 14-week-old male mice.^63^ In brief, after placing temporary ligatures around the proximal and distal aorta, the infrarenal aorta was infused with elastase from porcine pancreas (E1250, Sigma) for 5 min at 100mmHg. Body temperature was kept constantly at 37°C by a heating pad. After removing the infusion catheter, the aortotomy was sutured and the abdomen was closed. The abdominal segments were harvested at 3- or 28-days post-PPE surgery. For histological and immunofluorescence stainings the aorta was perfused with saline followed by 1.5% agarose before harvesting. For RNA isolation the aorta was collected in TRIzol™ Reagent (15596026, ThermoFischer scientific).

### Transmission electron microscopy

The abdominal aorta was harvested 28 days post PPE. The tissue was fixed in TEM fixative containing 2% glutaraldehyde and 2% formaldehyde in 0.1 M cacodylate buffer (pH 7.3) for 48h then washed with 0.1 M cacodylate buffer (pH 7.3). After washing, postfixation was applied using 2% OsO4 (Science Services) in 0.1 M cacodylate buffer for 2 h at 4°C followed by washing for four times with 0.1 M cacodylate buffer and dehydration in graded ethanol series (1×50%, 1×70%, 1×90%, 3×100%) for 15 min each. Sample were then incubated in a mix of 50% ethanol/propylenoxide then two times with pure propylenoxide for 15 min each. Samples were infiltrated with a mixture of 50% epon/propylenoxide and 75% epon/propylenoxide for 2 h each at 4°C and with pure Epon overnight at 4°C. The next day, epon was exchanged and samples were incubated for 2h at RT, placed into TAAB capsules and cured for 72 h at 60°C. Ultrathin (~D70 nm) sections were obtained using Leica EM UC6 Ultramicrotome (Leica, Germany) attached with diamond knife (DiATOME, Switzerland) and double-stained with 1.5% uranyl acetate (15 min at 37°C) followed by 3% lead citrate for 4 min. Images were acquired using a JEM-2100 Plus Transmission Electron Microscope (JEOL) operating at 80kV equipped with a OneView 4K camera (Gatan).

### Ultrasound imaging of the abdominal aorta

To obtain ultrasound images of the abdominal aorta, the mice were anesthetized with continuously delivered 2% isoflurane gas inhalation. Ultrasound gel was applied to the depilated skin of the abdomen and imaging was performed with a Vevo3100 imaging system (VisualSonics). B-mode, M-mode and EKV recordings were performed using a MX550D linear array transducer (25-55 MHz, Centre Transmit: 40 MHz, Axial Resolution: 40 μm) with a frame rate of 230–400 frames/s. All ultrasound images were analyzed using the Vevo 3100-software and parameters like aortic diameter, pulse wave velocity (PWV) and aortic wall thickenss were calculated.

### Immunofluorescence Staining

Abdominal aorta was harvested, cleaned and embedded into Tissue-Tek O.C.T. Compound (Sakura Finetek™ 4583) and stored at −80°C for cryopreservation. The tissue was cut into 10 μm thick sections using a Leica CM3050 S Cryostat (Leica, Germany). For Immunofluorescence stainings, the tissue sections were fixed with 4% paraformaldehyde (PFA) for 10 min and permeabilized using 0.1% Triton X-100 for 10 min at room temperature followed by blocking using 3% bovine serum albumin (BSA) for 1 h at room temperature. Tissue sections were incubated with primary antibody for anti-alpha smooth muscle Actin antibody (ab5694, Abcam) at 4 °C overnight. Normal rabbit IgG (Thermo Scientific, Waltham, MA, USA, 026102) was used for the negative controls. Next day, the sections were washed three times in PBS followed by incubation with Goat anti-Rabbit IgG (H+L) Cross-Adsorbed Secondary Antibody, Alexa Fluor™ 488 (A-11008, Invitrogen) and Alexa Fluor^®^ 594 anti-mouse CD68 Antibody (137020, BioLegend, USA) or Alexa Fluor^®^ 594 anti-mouse Ly-6G Antibody (127636, BioLegend, USA) for 1 h at room temperature in the dark followed by washing and mounting. Cell nuclei were stained with DAPI. Images were acquired with a BZ-X800 microscope system (Keyence, USA) and analyzed using ImageJ software.

### Histology

Masson Trichrome staining was performed using Trichrome Stain (Masson) Kit (HT15, Sigma) according to the manufacturer’s protocol. Images were acquired with a BZ-X800 microscope system (Keyence, USA) and analyzed using ImageJ software.

For elastin fibers, the tissue sections were imaged for the elastin autofluorescence using the GFP filter on BZ-X800 microscope system (Keyence, USA) and analyzed using ImageJ software. The processed images were graded as per the grading system outlined in Supplemental Figure S7A and S7B for MTS and elastin grading, respectively.^64^

### Flow cytometry analysis

Mice under 3% Isoflurane anesthesia were surgically opened, perfused with 10ml PBS (0.5 mM EDTA) via left ventricle. Abdominal aorta were isolated and digested for 1h, 37°C, 45rpm agitation in HBSS with 450U/ml Collagenase I (Sigma), 125U/ml Collagenase XI (Sigma) 60U/ml Hyalurase (Sigma), 60U/ml DNAse1 (ThermoFisher) and filtered through 100μm cell strainers (Sysmex) to achieve a single cell suspension. Samples were treated with FcR block (TruStain FcX, Biolegend, 1:100) and live-dead staining (Zombie UV, Biolegend) according to manufacturer’s protocol. Extracellular stainings were performed in FACS buffer, 20min, 4°C using the antibodies listed in the table below. Samples were acquired on an Aurora 5L Flow cytometer (Cytek Biosciences), unmixing and spillover correction was performed on SpectroFlo (Cytek Biosciences), conventional FCS gating and clustering was performed in Flowjo 10.8.1 with Phenograph and UMAP plugins. Samples of low cellularity were excluded from further analysis.

Immune cell populations were identified as CD45^+^, zombieUV^−^. Subpopulations were defined as Neutrophils (CD11b^+^, CD3^−^, Ly6G^+^); B-Cells (CD11b^−^, CD3^−^, CD19^+^, MHCII^+^); conventional T-cells (either CD11b^−^, CD3^+^, CD4^+^ or CD11b^−^, CD3^+^, CD8^+^); M1-macrophages (CD11b^+^, CD3^−^, F4/80^+^, MHCII^+^) or M2-macrophages (CD11b^+^, CD3^−^, F4/80^+^, MHCII^−^). Significant changes in cellular composition between the WT and *Mlkl^−/−^* were assessed by Welch ANOVA and Brown-Forsythe test in Prism9 (GraphPad Software). Abdominal aortic samples were further analyzed by UMAP and stratified according to surface marker expression.

| Antigen | Fluor | Clone | Dilution | Manufacturer |
| --- | --- | --- | --- | --- |
| CD45 | BUV496 | 30-F11 | 1:200 | BD |
| Ly6G | BUV563 | 1-A8 | 1:200 | BD |
| CD11b | BUV661 | M1/70 | 1:400 | BD |
| CD3e | BUV737 | 145-2C11 | 1:150 | BD |
| MeRTK | BV421 | 2B10C42 | 1:100 | Biolegend |
| CD11c | Pacific Blue | N418 | 1:200 | Biolegend |
| Ly6C | BV650 | HK1.4 | 1:100 | Biolegend |
| CD16.2 | BV711 | 9.00E+09 | 1:100 | Biolegend |
| CD8a | BV750 | 53-6.7 | 1:200 | Biolegend |
| CCR2 | BV786 | SA203G11 | 1:100 | Biolegend |
| CD115 | AF488 | AFS98 | 1:100 | Biolegend |
| CD4 | Spark Blue 550 | GK1.5 | 1:200 | Biolegend |
| TIMD4 | PE | RMT4-54 | 1:100 | Biolegend |
| F4/80 | PerCP/Cy5.5 | BM8 | 1:100 | Biolegend |
| Lyve-1 | PE/Cy7 | ALY-7 | 1:100 | Thermo |
| CD64 | APC | X54-5/7.1 | 1:100 | Biolegend |
| CD19 | Spark NIR 685 | 6D5 | 1:400 | Biolegend |
| MHCII | AF700 | M5/114.15.2 | 1:100 | Biolegend |
| CX3CR1 | APC/Fire750 | SA011F11 | 1:200 | Biolegend |

### mRNAseq analysis

Total RNA was extracted from cleaned abdominal aortas 3 days post PPE from WT, *Ripk1*^*D138N/D138N*^, *Mlkl*^*AA*^ and *Mlkl^−/−^* animals (n = 4 each), abdominal aortae from untreated WT (WT baseline) animals were used as control. RNA was shipped to Novogene (Cambridge, UK) where the mRNA sequencing was performed as per their in-house protocols outlined in Supplementary Method Section. Gene expression read counts were exported and analyzed using the integrated Differential Expression and Pathway (iDEP) tool (http://bioinformatics.sdstate.edu/idep96/) to identify the differentially expressed genes (DEGs) and enriched GO terms, results were visualized with GOplot package^65^ volcano plots are created using VolcaNoseR web application^66^.

### Bone marrow transplantation

In a bone marrow (BM) transplantation study irradiated recipient WT mice were reconstituted with *Mlkl*^*AA*^-BM (WT^MLKLAA-BM^) and irradiated *Mlkl*^*AA*^ recipient mice were reconstituted with WT-BM (MLKL^AA+WT-BM^) resulting in SMC (*WT*)/ macrophage (*Mlkl*^*AA*^) and SMC (*Mlkl*^*AA*^)/ macrophage (WT) chimera mice. Mice were given neomycin (1.6 mg/ml) through drinking water 8 days before the irradiation. Recipient mice were irradiated with□10□Gy whole-body irradiation in a Cesium-137 gamma source (Biobeam GM 8000). Bone marrow cells from the donor animals were isolated from the femurs and resuspended in cold PBS and 5 × 10^6^ bone marrow cells (150 μL) were injected by tail vein into each of the recipient mice 24h after irradiation. Four weeks after transplantation, small amount of blood was withdrawn and platelet counts were determined by using Element HT5 (scil animal care company GmbH, Germany) and animals were subjected to PPE surgery. Transplantation efficacy was determined by flow cytometry for CD45.1 (donor) and CD45.2 (recipient) 5 days before PPE and reached 95% cell reconstitution in all animals.

## In vitro methods

### Reagents

All the cell culture reagents such as DMEM, RPMI, puromycin, TrypLE™ select enzyme, FBS, HBSS buffer, etc. were purchased from Gibco/Thermo Scientific (Waltham, MA, USA). Necrosulfonamide was purchased from Tocris (Cat. No. 5025/10). Elastase from porcine pancreas was purchased from Sigma (Cat. No. E1250). Doxycycline Hydrochloride as purchased from Sigma (Cat. No. D3072-1ML).

### Cell culture

Human aortic SMC (HAoSMC) were purchased from ATCC (PCS-100-012, ATCC) and propagated in Vascular Cell Basal Medium (PCS-100-030, ATCC) supplemented with Vascular Smooth Muscle Cell Growth Kit (PCS-100-042, ATCC) as per the manufactures protocol in standard cell culture incubator at 5% CO_2_, 37 °C.

HL-60 cells were maintained in RPMI-1640 (72400047, Gibco) with 10% FBS (F9665, Sigma), 1x GlutaMAX™ Supplement (35050061, Gibco), 1x Penicillin-streptomycin (15140122, Gibco) in a 5% CO_2_ atmosphere at 37 °C. The HL-60 cells were differentiated in to a neutrophil-like (neut-HL60) phenotype by culturing in a RPMI-1640 medium supplemented with 10% FBS and 1.3% DMSO for 6 days.^67^ Presence of neutrophil-like cells was confirmed by staining the cells with Ly6G and DAPI.

### MLKL cloning and generation of stable SMC lines

To induce necroptosis in SMCs we opted for inducible MLKL overexpression system. The full-length MLKL and N-domain MLKL (amino acid 1 – 182) were amplified from cDNA clone (RC213152, Origene, USA) using two step PCR reaction with primers mentioned in the table below. The primers are designed to incorporate the attL1 and attL2 overhangs on either side of the PCR product to facilitate the ligation using LR-Clonase enzyme. The PCR product was then ligated into pB-TAC-ERP2 plasmid (PB-TAC-ERP2 was a gift from Knut Woltjen (Addgene plasmid # 80478)) and positive clones were confirmed by DNA sequence analysis.

To generate the HAoSMC lines overexpressing hMLKL, the cells were transfected with donor pB-TAC-ERP2 vector containing either full-length hMLKL (F-hMLKL) or N-domain hMLKL (ND-hMLKL) isoform along with pCAGPBase plasmid (pCAGPBase was a gift from Joseph Loturco (Addgene plasmid # 40972)) encoding for transposase. 48 h post transfection the cells were selected using 2 μg/ml of puromycin. Post transfection the serum in the growth media was replaced with Tet system approved serum (A4736101, ThermoFischer). Cells were treated with 1 μg/ml Doxycycline to induce the MLKL overexpression and for all experiments the cells between passage three and seven were used.

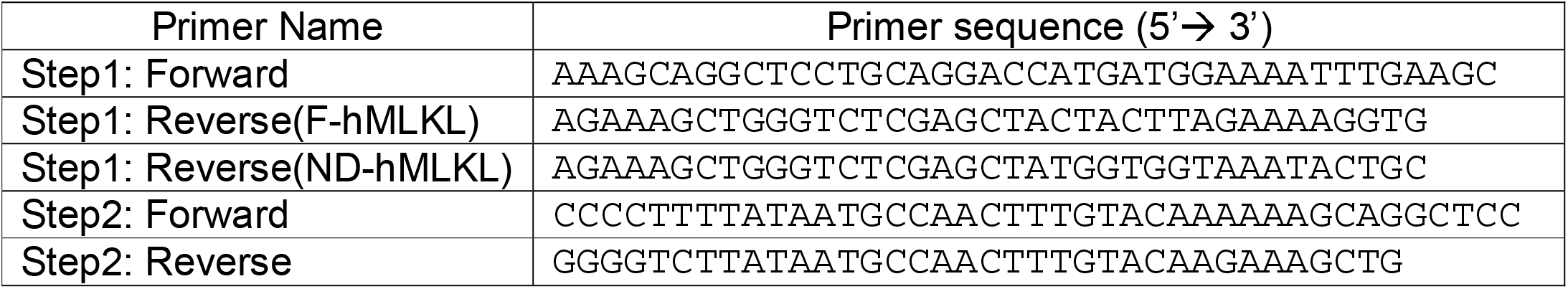

### Cytokine Array

A Human Cytokine Antibody Array (ab133998, Abcam) was used to analyze the supernatant of hSMCs as per the manufacturer’s instructions. In brief, 0.1×10^6^ hSMCs were seeded in 6 well plate and cultured till cells reached ~70% confluence. Then doxycycline was added and supernatants were collected after 24h for the measurements of released cytokines. SMCs without doxycycline were used as uninduced control. The dot blots were used to calculated the integrated densities using ImageJ and ‘Protein Array Analyzer’ Tool.^68^

### Statistical Analysis

Data are presented as mean ± SEM. Shapiro–Wilk test and Brown–Forsythe test were utilized to test for normal distribution and equality of variances, respectively. Differences between three or four groups were evaluated using one-way or two-way repeated measures analysis of variance (ANOVA) with post-hoc Tukey’s test if data were normally distributed and variances were equal. Kruskal–Wallis test with Dunn’s post-hoc test was performed if data were not normally distributed. For comparison of two data sets consisting of normally distributed data, the unpaired t-test was utilized. A P-value <0.05 was considered statistically significant. All statistical analyses were performed using GraphPad Prism 8.4.0 (GraphPad Software, San Diego, CA, USA).

## Author Contributions

MM, MA and MP conceived and supervised the study. HNN harvested the animals, gathered and analyzed data, and wrote the manuscript. DM, HL, ZY, PS performed echocardiographic, histo- and IF-stainings. FP and HW performed FACS and analyzed the data, SG performed the tail vein injections for bone marrow experiments, GD, HG, AH, FSN, AS, AGS, SB performed experimental support, MM provided support with mouse lines, surgeries and ethics applications. All authors participated in data analysis, critically reviewed the manuscript, and gave final approval.

## Acknowledgments

The authors would like to thank Christina Vosen, Sharon Weingarten, Katharina Tinaz, Nadja Klein and Simon Grimm for excellent technical support. We also thank the CECAD Imaging Facility, Dr. Astrid Schauss and Ms. Beatrix Martiny for their support in transmission electron microscopy.

## Figure Legends

**Supplemental Figure S1**

(**A**) Occurrence of aortic dilation up to 28 days after PPE in the indicated genotypes.

(**B**) Representative transmission electron microscopy (TEM) images of abdominal aortic sections at baseline and 28 days post AAA. Scale bar = 1000nm.

**Supplemental Figure S2**

Lack of MLKL reduces infiltration of the proinflammatory immune cells. Representative confocal images of aSMA (green) and Ly6G (red, left) and aSMA (green) and CD68 (red, right) of (**A**) WT baseline aortae and (**B**) aortae 28 post PPE of the indicated genotypes (scale bars = 100μm and 20μm; Lumen (L), media (M), adventitia (Ad)). (**C**) Quantification of total aortic number of CD68^+^ cells and CD68^+^ cells in the aortic media of day 28 post-PPE. Statistical significance was determined by one-way ANOVA with Tukey’s multiple comparisons test. * = p<0.05, ** = p<0.01, *** = p<0.005, ns = not significant. (**D**) Pregating strategy to determine blood and aortic PMN. (**E**) Cell clusters and heatmap of cell specific marker expression analyzed by FACS.

**Supplemental Figure S3**

(**A and B**) GOcircle plot display upregulated and downregulated genes from WT-d3-PPE animals with their log fold change (logFC) and the most statistically significant GO terms (table) respectively. (**C**) Genes associated with fibrinolysis pathway with their fold change values from animals after 3 days of PPE. (**D**) Genes only downregulated in Mlkl^−/−^ d3 PPE compared to WT d3 PPE. (**E, F and G**) Heat map of DEGs from *Mlkl^−/−^* d3 PPE, *Mlkl*^*AA*^ d3 PPE and *Ripk1*^*D138N/D138N*^ d3 PPE animals respectively.

**Supplemental Figure S4**

Chord plot illustrating the most significant GO biological process terms and the genes contributing to that enrichment (Left) arranged in order of their logFC expression level. (**A**) enrichment analyses for DEGs from *Mlkl^−/−^* d3 PPE animals. Downregulated (left) and upregulated (right), (**B**) enrichment analyses for DEGs from *Mlkl*^*AA*^ d3 PPE animals. Downregulated (left) and upregulated (right), (**C**) enrichment analyses for DEGs from *Ripk1*^*D138N/D138N*^ d3 PPE animals. Downregulated (left) and upregulated (right).

**Supplemental Figure S5**

Loss of MLKL in aortic SMCs is protective against AAA. (**A**) Experimental scheme of the bone marrow transplantation experiment. (**B**) Representative macroscopic images of complete abdominal aortic rings stained with Masson’s Trichrome Staining (MTS) (top panel) and elastin autofluorescence (green; bottom panel). (**C**) Representative confocal images of complete abdominal aortic rings with aSMA (green) and Ly6G (red; top panel) and CD68 (red; bottom panel) (scale bars = 100 μm).

**Supplemental Figure S6**

Inhibition of MLKL reduces release of proinflammatory cytokines. (**A**) Quantification of cytokine array dot blots of untreated and induced ND-MLKL-hSMCs using ‘Protein Array Analysis’ tool in ImageJ. The integrated densities were calculated and normalized with respective integrated positive controls (POS) on each blot and then compared with untreated samples.

**Supplemental Figure S7**

Definition and representative images of (**A**) Masson’s Trichrome Staining (MTS) and (**B**) elastin degradation grade.

**Supplemental Video V1**

Dox treated ND-MLKL-SMC cocultured with dHL60 and live cell imaging. The video shows typical necroptotic cell death with initial rounding of the cells followed by membrane rupture in the SMCs upon overexpression of MLKL which was monitored indirectly via mCherry expression.

**Supplemental Video V2**

Dox treated ND-MLKL-SMC cocultured with dHL60 and live cell imaging was performed. The video shows dHL60 cells interacting with necroptotic SMC.

