## Supplementary figures and images for "Inhibition of MLKL impairs abdominal aortic aneurysm development by attenuating smooth muscle cell necroptosis"

### Supplemental Figure S1

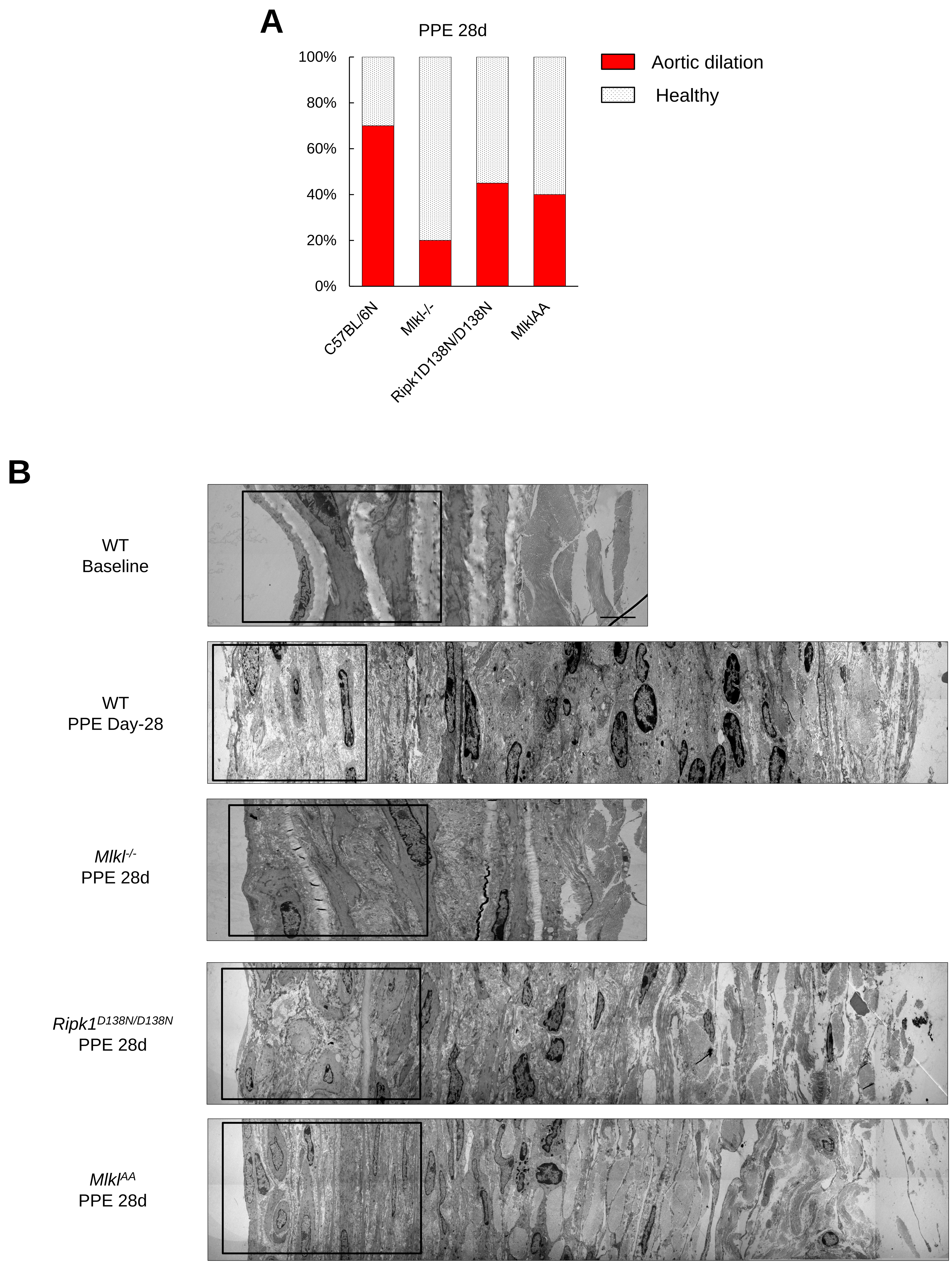

### Supplemental Figure S2

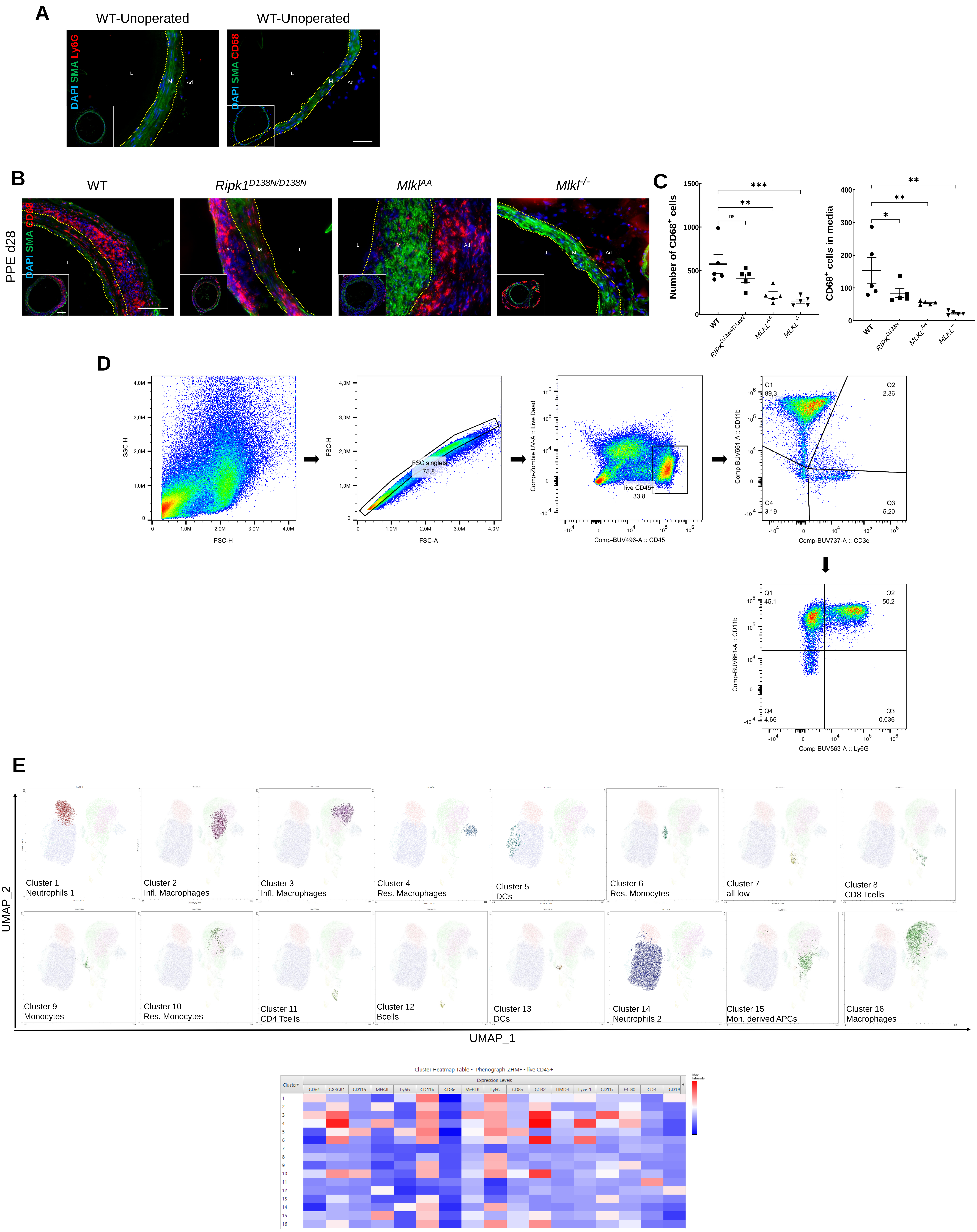

### Supplemental Figure S4

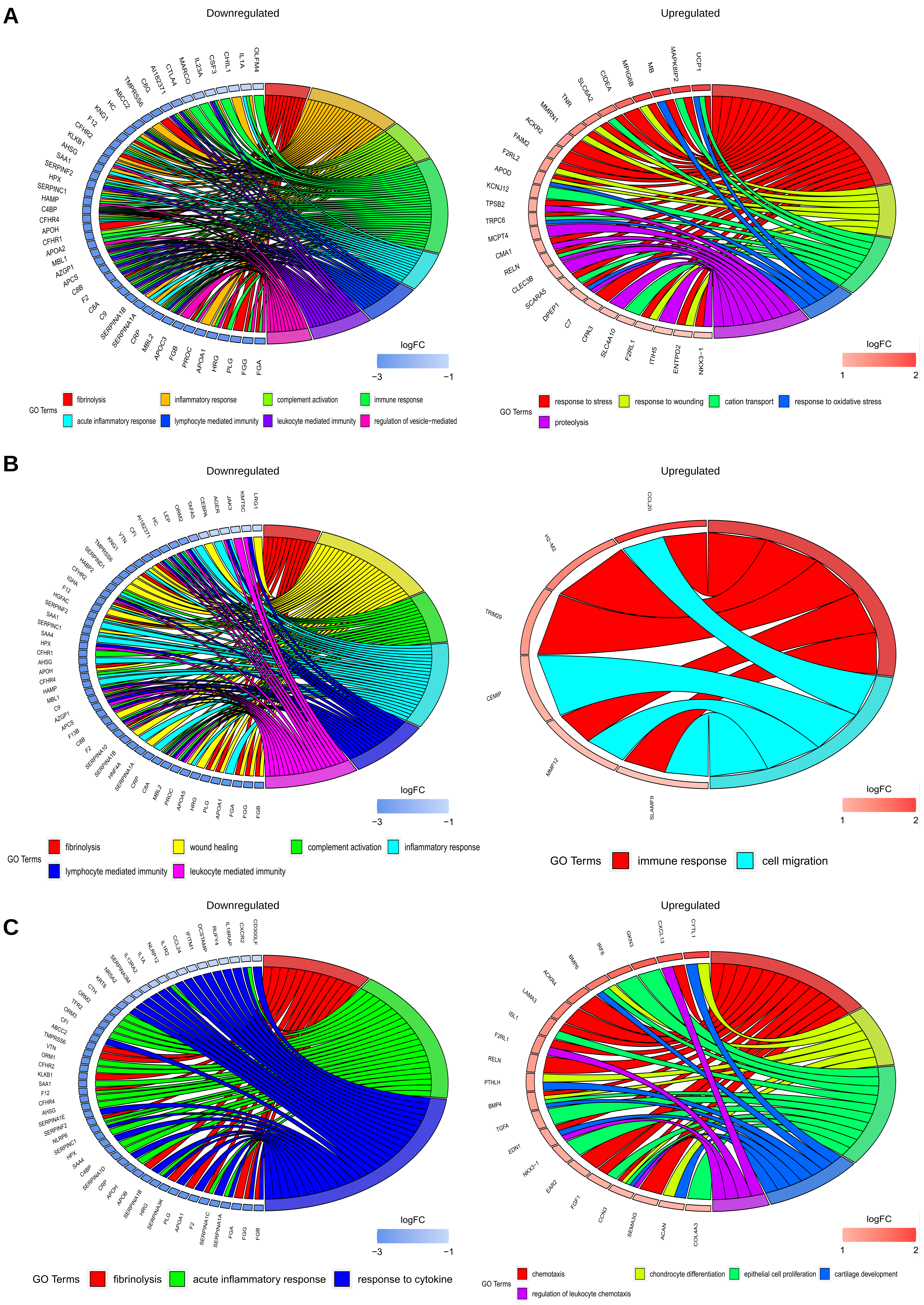

### Supplemental Figure S5

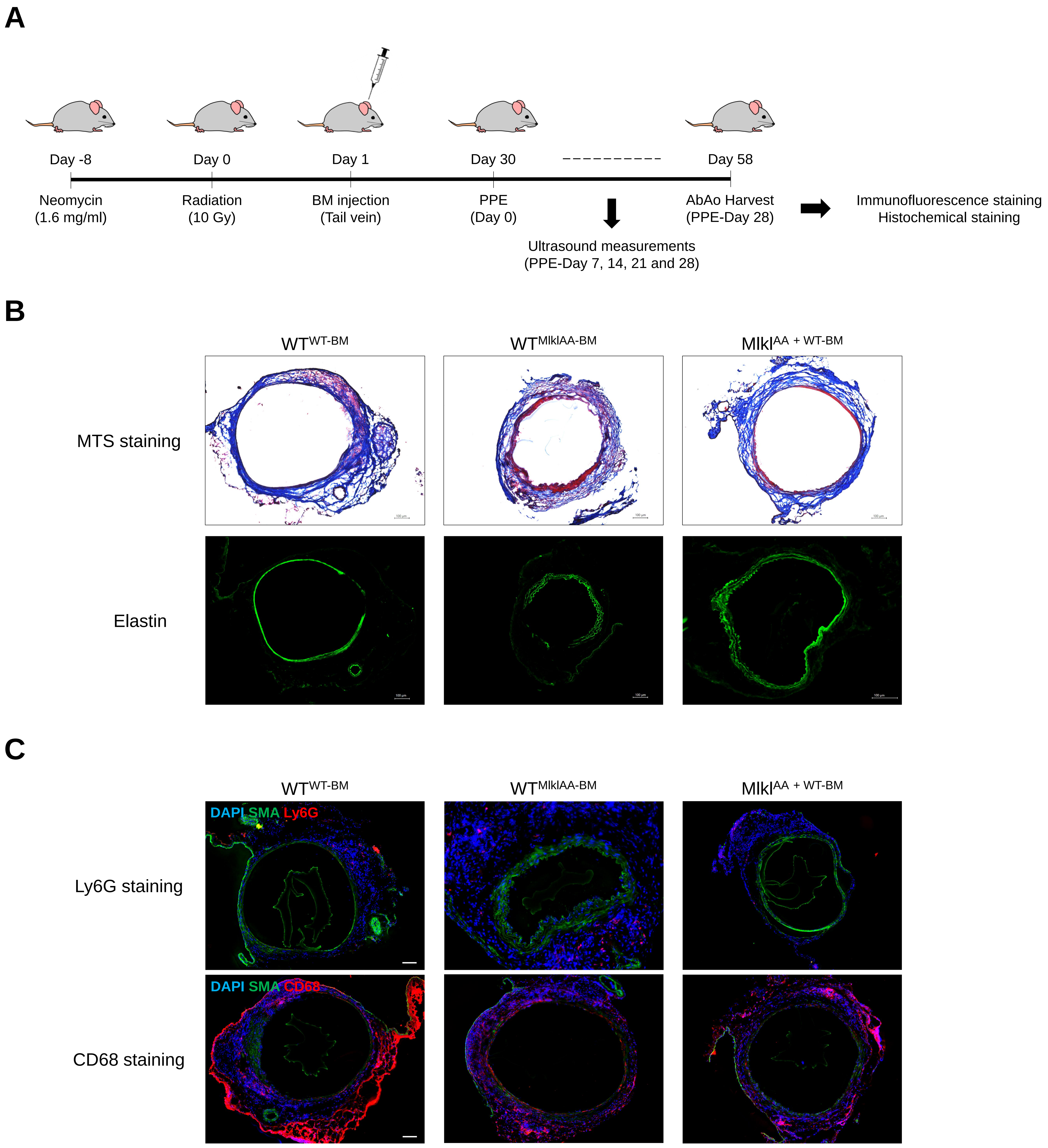

### Supplemental Figure S6

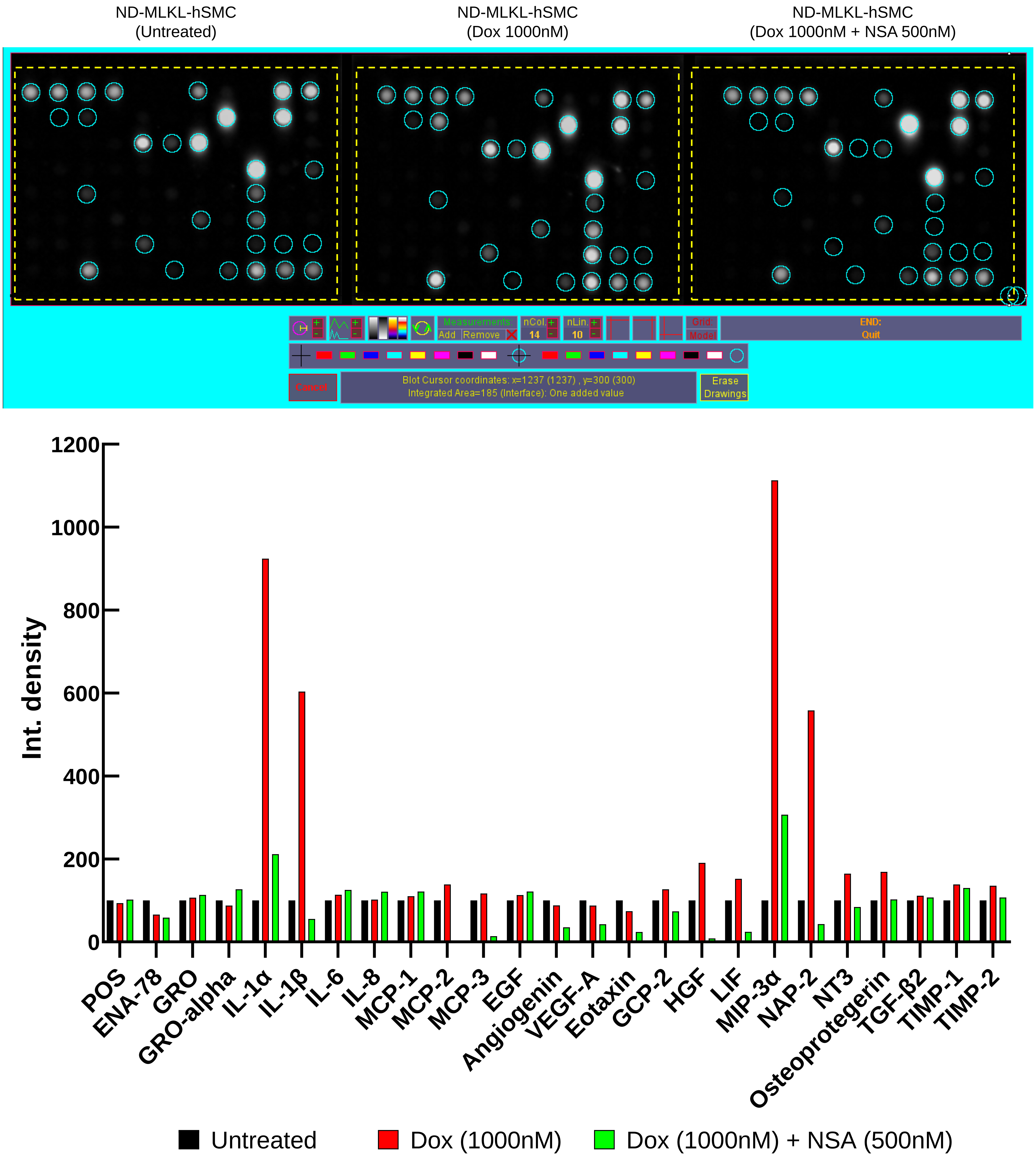
