## Supplemental Figure S3 for "Inhibition of MLKL impairs abdominal aortic aneurysm development by attenuating smooth muscle cell necroptosis"

**A**

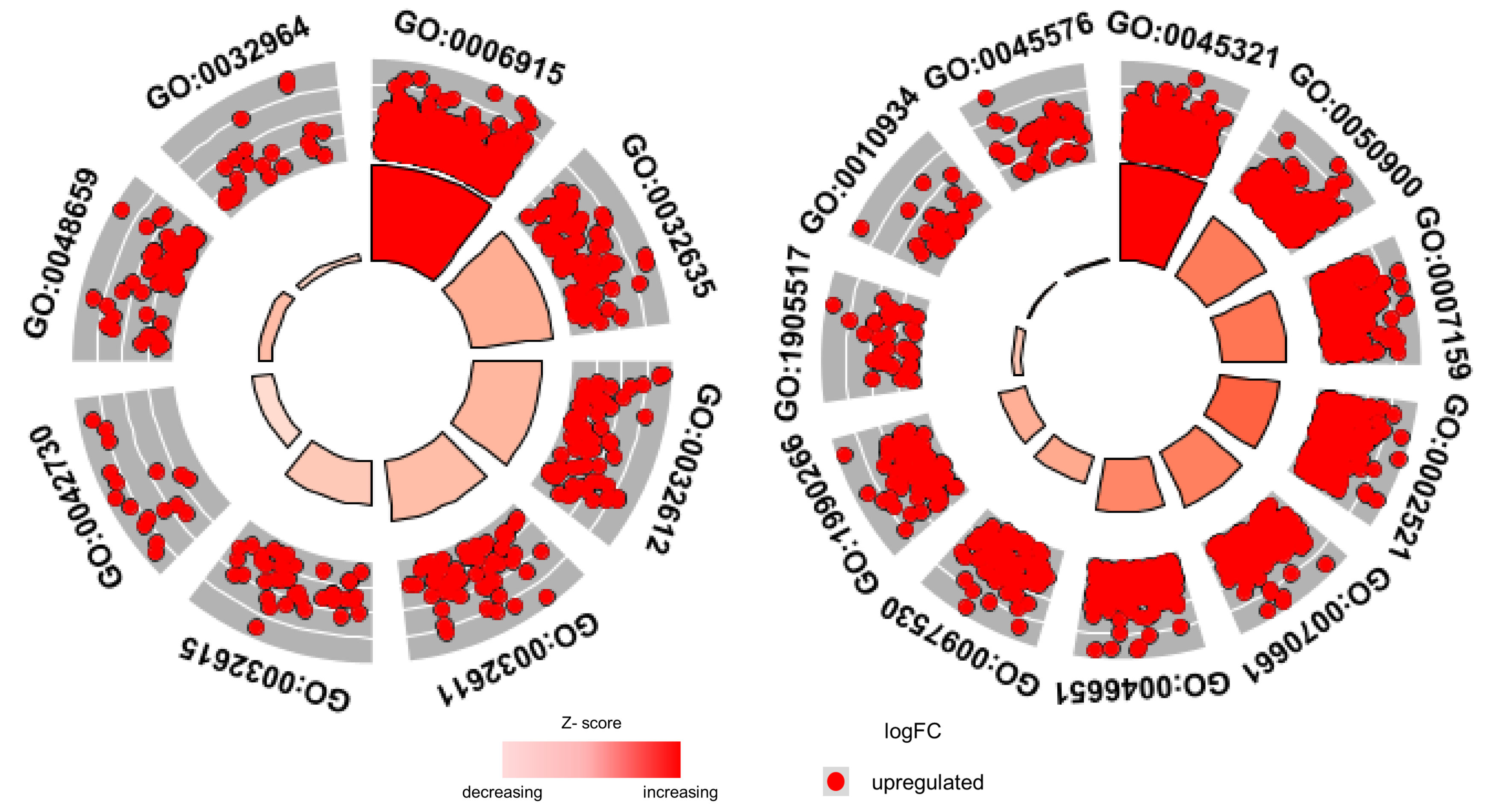


| ID | Description |
|---|---|
| GO:0006915 | apoptotic process |
| GO:0032635 | interleukin-6 production |
| GO:0032612 | interleukin-1 production |
| GO:0032611 | interleukin-1 beta production |
| GO:0032615 | interleukin-12 production |
| GO:0042730 | fibrinolysis |
| GO:0048659 | smooth muscle cell proliferation |
| GO:0032964 | collagen biosynthetic process |

| ID | Description |
|---|---|
| GO:0045321 | leukocyte activation |
| GO:0050900 | leukocyte migration |
| GO:0007159 | leukocyte cell-cell adhesion |
| GO:0002521 | leukocyte differentiation |
| GO:0070661 | leukocyte proliferation |
| GO:0046651 | lymphocyte proliferation |
| GO:0097530 | granulocyte migration |
| GO:1990266 | neutrophil migration |
| GO:1905517 | macrophage migration |
| GO:0010934 | macrophage cytokine production |
| GO:0045576 | mast cell activation |

**B**

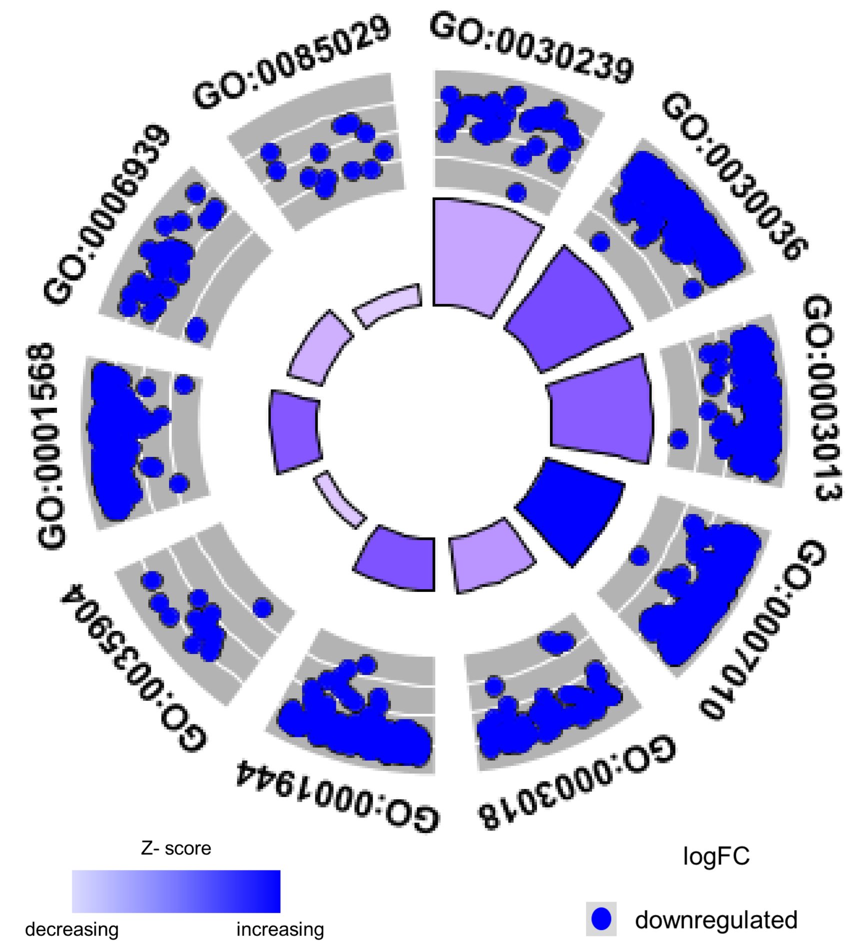


| ID | Description |
|---|---|
| GO:0030239 | myofibril assembly |
| GO:0030036 | actin cytoskeleton organization |
| GO:0003013 | circulatory system process |
| GO:0007010 | cytoskeleton organization |
| GO:0003018 | vascular process in circulatory system |
| GO:0001944 | vasculature development |
| GO:0035904 | aorta development |
| GO:0001568 | blood vessel development |
| GO:0006939 | smooth muscle contraction |
| GO:0085029 | extracellular matrix assembly |

**C**

**Fibrinolysis associated genes**

| Gene Name | Gene ID | logFC | | | |
|---|---|---|---|---|---|
| | | WT-D3-PPE | Mlkl$^{-/-}$-D3-PPE | Mlkl$^{AA}$-D3-PPE | Ripk1$^{-D138N/D138N}$-D3-PPE |
| apolipoprotein H | Apoh | 4.45 | -5.87 | -5.68 | -7.09 |
| coagulation factor II | F2 | 4.86 | -7.72 | -7.23 | -8.91 |
| fibrinogen alpha chain | Fga | 5.05 | -10.7 | -11.41 | -9.86 |
| fibrinogen beta chain | Fgb | 4.82 | -8.7 | -11.73 | -10.18 |
| fibrinogen gamma chain | Fgg | 4.97 | -8.98 | -10.74 | -11.33 |
| histidine-rich glycoprotein | Hrg | 5.32 | - | - | -8.98 |
| plasminogen | Plg | 4.43 | -8.77 | - | -9.38 |

**D**

| Gene Name | Gene ID | logFC |
|---|---|---|
| glycine-N-acyltransferase | Glyat | -6.57 |
| mixed lineage kinase domain-like | Mlkl | -2.24 |
| cellular retinoic acid binding protein I | Crabp1 | -1.82 |
| cytotoxic T-lymphocyte-associated protein 4 | Ctla4 | -1.75 |
| macrophage receptor with collagenous structure | Marco | -1.75 |
| colony stimulating factor 3 (granulocyte) | Csf3 | -1.23 |
| chitinase-like 1 | Chil1 | -1.20 |
| predicted gene 14548 | Gm14548 | -1.20 |
| ribosomal protein L30, pseudogene 10 | Rpl30-ps10 | -1.10 |
| olfactomedin 4 | Olfm4 | -1.05 |
| predicted gene 12191 | Gm12191 | -1.05 |

**E**

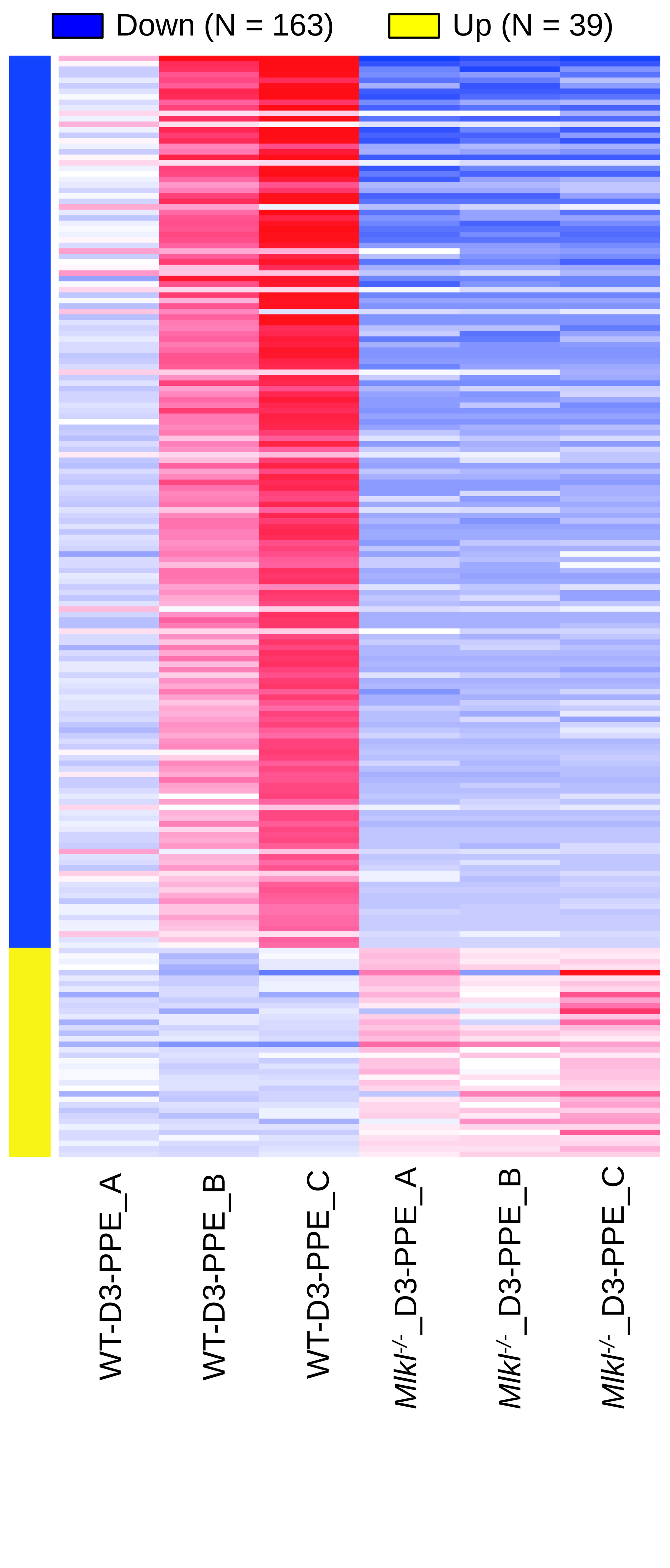


**F**

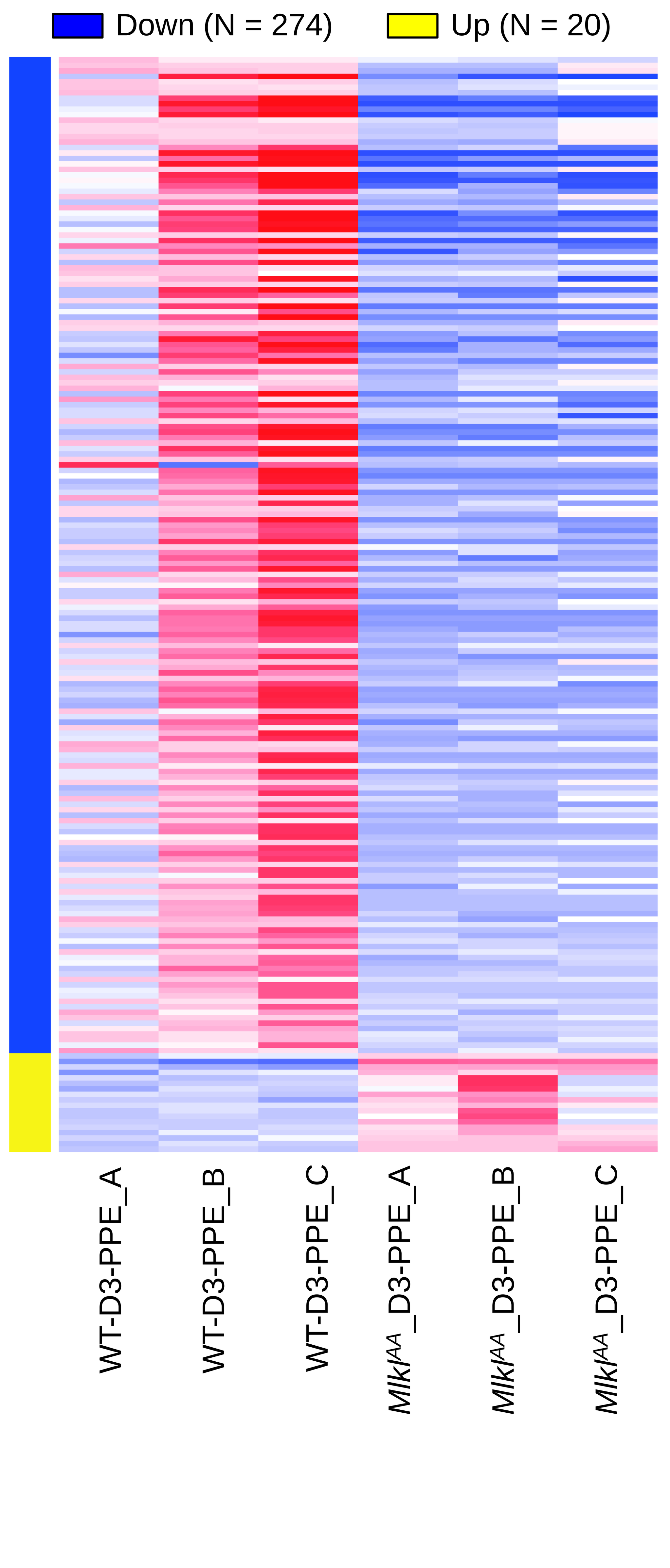


**G**

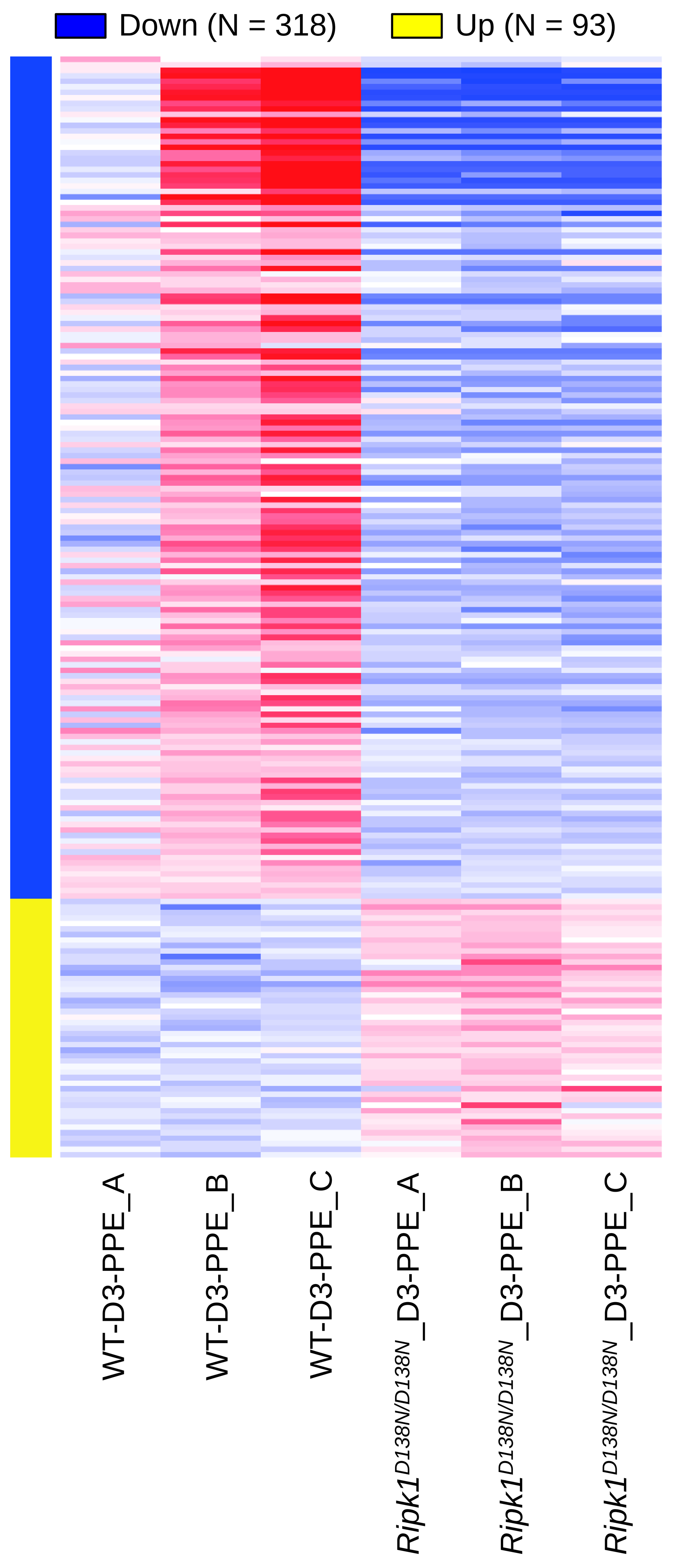
