## Supplemental Figure S7 for "Inhibition of MLKL impairs abdominal aortic aneurysm development by attenuating smooth muscle cell necroptosis"

A

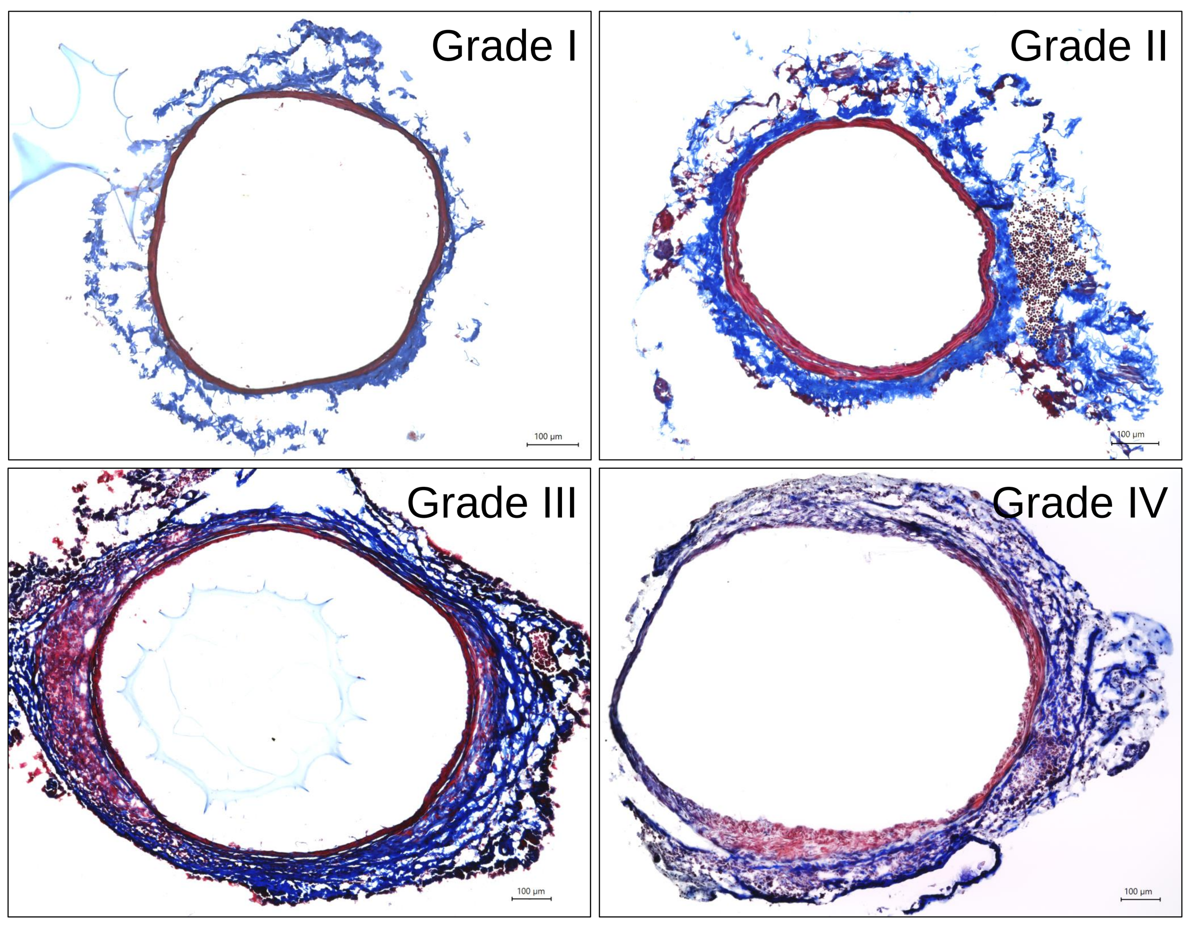


| MTS grading | |
|---|---|
| Grade I | Intact media, wavy collagen fibers bundles (Unoperated (Control) aorta) |
| Grade II | Small increase in the media content, loss of waviness in collagen fibers bundles |
| Grade III | Significant increase in the media content and loss of media boundary, loss of waviness in collagen fibers bundles |
| Grade IV | Significant increase in the media content and loss/ rupture of media boundary, loss of waviness in collagen fibers bundles |

B

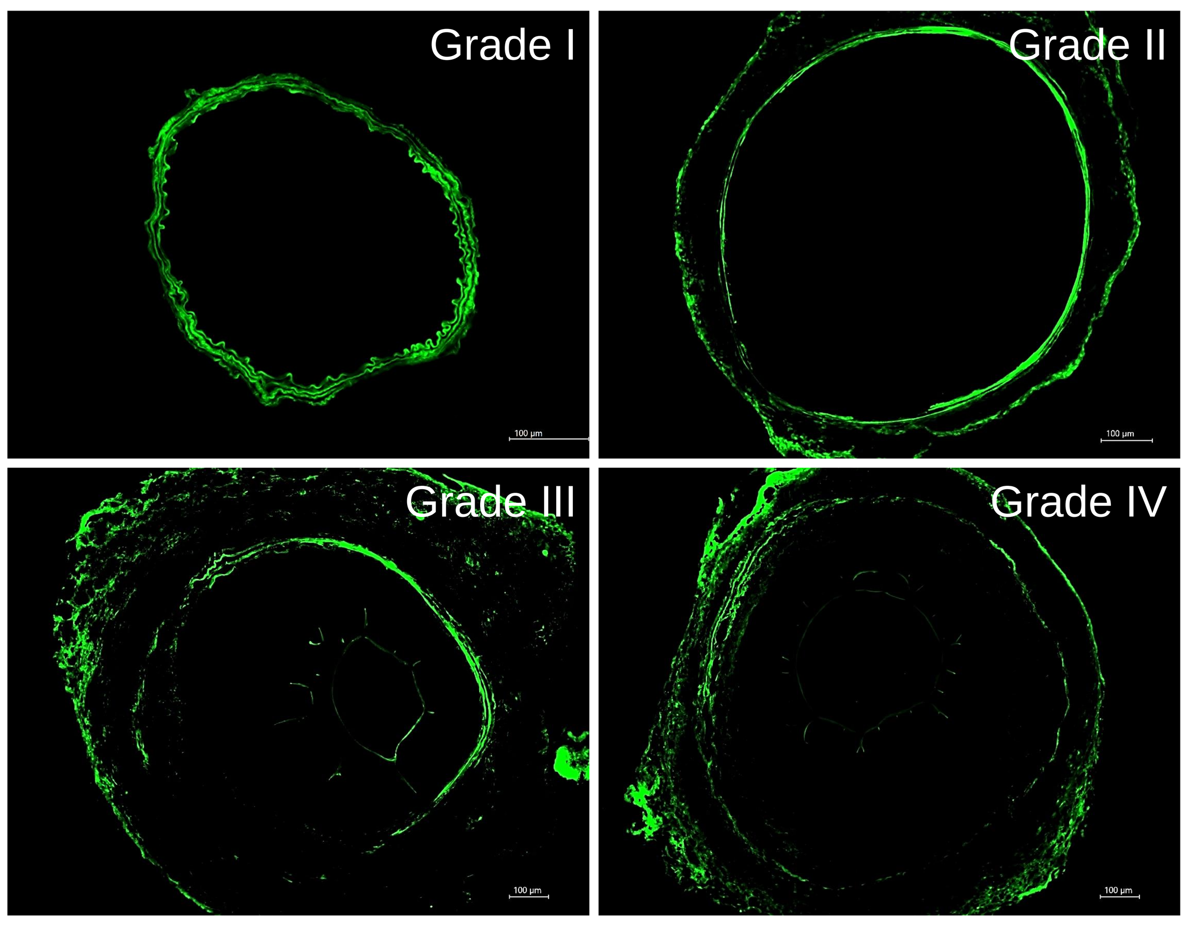


| Elastin degradation grading | |
|---|---|
| Grade I | Intact/ wavy elastin fibers (Unoperated (Control) aorta) |
| Grade II | Visible elastin breaks, loss of waviness in elastin morphology |
| Grade III | Visible elastin breaks, loss of ~50% of elastin signal |
| Grade IV | Visible elastin breaks, loss of >50% of elastin signal |
